# Oocyte polarity is established independently of the Balbiani body

**DOI:** 10.1101/2025.08.26.672409

**Authors:** Manami Kobayashi, Joseph Zinski, Mary C. Mullins

## Abstract

In most animals, oocyte polarity establishes the embryonic body plan by asymmetrically localizing axis-determining transcripts. These transcripts first localize in *Xenopus* and zebrafish oocytes to the Balbiani body (Bb), a large membrane-less organelle conserved from insects to humans. The Bb is transient, disassembling and anchoring at one pole the axis-determining transcripts that establish the vegetal pole of the oocyte. Aggregation of the Bb depends on the Bucky ball (Buc) protein, an intrinsically disordered protein with a prion-like self-aggregation domain. In zebrafish *buc* null mutants, the Bb fails to form and oocytes lack polarity. Here, we established *buc* hypomorphic mutants that fail to form the Bb, but remarkably Buc protein and vegetal mRNAs localized normally at the vegetal cortex of the oocyte. Thus, these *buc* hypomorphic mutants displayed normal oocyte polarity, demonstrating that the Bb is not required to establish oocyte polarity. We found that both a reduced Buc protein level and truncation of the N-terminal 10 amino acids contribute to Bb failure in the hypomorphic mutants.

## INTROUDCTION

Cell polarity establishes specialized functions to opposing regions of a cell. Alterations in cell polarity can lead to morphogenic abnormalities of a tissue and diseases like cancer. Cell polarity is a hallmark of embryogenesis that is evident in the fertilized egg along the animal-vegetal (A-V) axis. The animal pole is defined as the region where the polar bodies are extruded, whereas the vegetal pole is opposite to it. In most animals, polarity of the egg is established during oogenesis, where patterning cues that are localized along the A-V axis subsequently define the dorsal-ventral (D-V), anterior-posterior, and left-right axes of the embryo^1–3^. In Xenopus and zebrafish, body-axis-formation related mRNAs are localized at the vegetal cortex of the oocyte and egg^4–8^. Mis-localization or loss of these body-axis-formation related mRNAs causes severe ventralization of the body axis^6–9^. However, how oocyte polarity is established during oogenesis is not fully understood.

The first polarized structure of the oocyte is the Balbiani body (Bb), which is also known as the mitochondrial cloud in Xenopus. The Bb is a large cytoplasmic membrane-less organelle observed in the primary oocyte, which contains mitochondria, Golgi apparatus, endoplasmic reticulum and RNA-protein complexes^10^. This structure is conserved from insects to mammals, including in humans^10–16^. Studies in *Xenopus* oocytes indicate that the Bb facilitates the translocation of vegetal mRNAs to the vegetal cortex during oogenesis^17^. For example, vegetal mRNA *Xlsirts*, *Xcat2/nanos* and *Xwnt11* localize to the Bb, which then translocate to the oocyte cortex, where the Bb disassembles, leaving these mRNAs anchored at the vegetal cortex^17^. The Bb initially forms at the periphery of the oocyte nucleus, grows and begins to disperse at the future vegetal cortex in later stage oocytes^10,18^. Thus, the Bb is postulated to specify the vegetal pole of the oocyte during oogenesis.

In a zebrafish maternal-effect mutant screen, we identified *bucky ball* (*buc*) as a key regulator of Bb and A-V polarity formation^19,20^. In *buc* null mutant oocytes, the Bb fails to form, and mRNAs normally localized at the vegetal pole are unlocalized, while animally-localized mRNAs are radially expanded around the oocyte cortex^20^. Embryos from *buc* null mutant females are similarly affected^19^. Buc is a highly disordered protein that can form amyloid-like self-aggregates via its prion-like domain^21^. This property of Buc protein is postulated to contribute as a critical condensation factor of the Bb. Consistent with the model in *Xenopus* oocytes, these results suggest that the absence of the Bb leads to the failure of A-V polarity establishment in *buc* null mutants. However, in this context, it is unclear whether the Bb itself and/or Buc itself is important for A-V axis formation.

Here, we established using CRISPR/Cas9 genome editing in zebrafish, *buc* hypomorphic mutant lines that differ to *buc* null alleles. The start codon is mutated in *buc* hypomorphic mutants, causing an N-terminal truncation of the Buc protein. *buc* hypomorphic mutant females produced embryos with a normal A-V axis but with a fraction displaying a ventralized phenotype. Surprisingly, despite the normal A-V embryonic axis, the Bb failed to form in stage I oocytes of the *buc* hypomorphic mutants. In stage II oocytes, remarkably, we found Buc protein and localized mRNAs at the vegetal cortex in the absence of Bb formation, with the weaker allele displaying normal oocyte A-V polarity. These results demonstrate that the Bb is not required for A-V polarity formation, but Buc itself plays a key role to establish oocyte polarity. Then, we addressed whether reduced Buc protein levels could be the cause for failed Bb formation in *buc* hypomorphic mutants. Quantitative western blots showed a significant reduction in Buc protein levels in the mutants. Furthermore, we established a transgenic line that overexpresses the mutant Buc protein in the *buc* hypomorphic background. In some of these transgenic oocytes, we observed a nearly normal Bb, but a large fraction also exhibited an abnormal Bb or Buc aggregates. These results indicate that a reduced protein amount and N-terminal truncation of Buc cause the Bb phenotype. Moreover, in the strong hypomorphic allele, we found that the *dazl* mRNA localization domain was reduced vegetally, whereas the animal pole mRNA *cyclin B1* domain was expanded and the Buc protein expression level was greatly reduced. These results provide insights into the function of Buc in oocyte polarity formation and reveal a Bb-independent mechanism that establishes oocyte polarity.

## RESULTS

### *buc* hypomorphic alleles display a normal A-V axis but are severely ventralized

Using the CRISPR/Cas9 system, we established new *bucky ball* (*buc*) mutant lines, *buc^p6del^* and *buc^p6del11in^*(hereafter, we will refer to *buc^p6del^*as *buc^Δ6^* and *buc^p6del11in^*as *buc^Δ6+11^*), altering the start codon: 6 bp are deleted in *buc^Δ6^* and 6 bp are deleted and 11 bp are inserted in *buc^Δ6+11^* (Fig. 1A). We performed RT-PCR using total RNA from each mutant allele and confirmed by sequencing that 6 nucleotides are deleted of the *buc^Δ6^* transcript that comprise the first two codons of the open reading frame (Fig 1A). For the *buc^Δ6+11^* transcript, we found that although the splice site is intact, it skips exon 2, resulting in deletion of 7 nucleotides of the coding sequence and 51 nucleotides of the 5’UTR (Fig. 1A-B, S1A). For both mutant transcripts, the next in-frame start codon is in exon 3, so we predict that these mutations cause a 10-amino acid N-terminal truncation of the Buc protein (Fig.1C).

**Figure 1.**
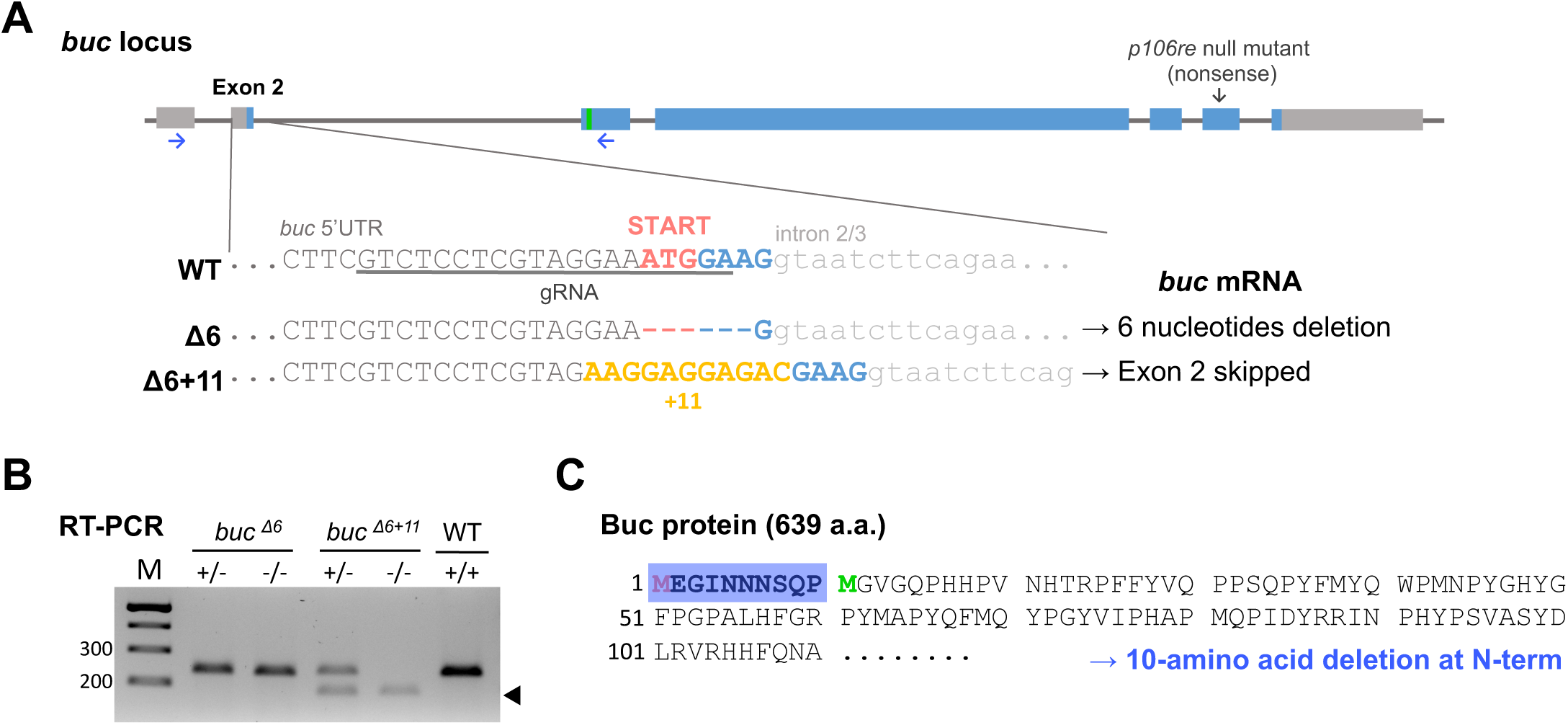
*buc* CRISPR mutants have predicted 10-amino acid N-terminal truncation of Buc protein. **(A)** Generating *buc* CRISPR mutant lines. CRISPR guide RNA was designed to the start codon in exon 2 (underline). 6 bp are deleted in the *buc^Δ6^* allele and a 6 bp deletion and 11 bp insertion were found in the *buc^Δ6+11^* allele. Green box in exon 3 indicates next in-frame ATG. Black arrow in exon 6 indicates a nonsense mutation in the *buc^p106re^*null mutant. **(B)** RT-PCR of maternal *buc* mRNA from the CRISPR mutants. cDNA was prepared from unfertilized eggs, and the primers designed to exon 1 and exon 3 (blue arrows in panel A) were used for RT-PCR. Black arrowhead indicates exon 2 skipped products in *buc^Δ6+11^*. See Figure S1A for *buc* mutant sequencing results. M is bp markers. **(C)** Predicted mutant Buc protein. Both *buc* CRISPR mutants lack the start codon in exon 2 and the next in-frame methionine (green M, predicted alternative start site) is in exon 3. Thus a 10-amino acid deletion at the N-terminus of the Buc protein (blue box) is predicted in both mutants. See Figure S1B for *buc* mRNA sequence.

Next, we examined the maternal-effect embryonic phenotype of the *buc* CRISPR alleles. Zebrafish *buc^p106re^*null mutant females produce embryos that lack animal-vegetal (A-V) polarity^19,20,22,23^, displaying radially distributed cytoplasm and multiple micropyles at the one-cell stage, and dying before 1-day post fertilization (dpf) (Fig. 2A, S2A-B). On the other hand, embryos from *buc^Δ6^* and *buc^Δ6+11^* homozygous females showed a normal A-V axis and a single micropyle typically, and could develop to adulthood (Fig. 2A, S2A-B). However, some embryos displayed a severely ventralized phenotype (Fig. 2A-B). These phenotypes were not observed in embryos from heterozygous females crossed to homozygous males, so *buc^Δ6^* and *buc^Δ6+11^* are recessive maternal-effect mutant alleles (hereafter ‘mutant’ refers to a homozygous female unless otherwise described). The ventralized rate and phenotypic severity varied from female to female and cross to cross (Fig. S2C-D). We found that *buc^Δ6^* mutant females that produced the strongest ventralized phenotype were similar to the mild *buc^Δ6+11^* female phenotypic group. Overall, *buc^Δ6+11^* showed a more severe ventralized phenotype than *buc^Δ6^* and both alleles represent hypomorphic alleles of the *buc* gene (Fig. 2B).

**Figure 2.**
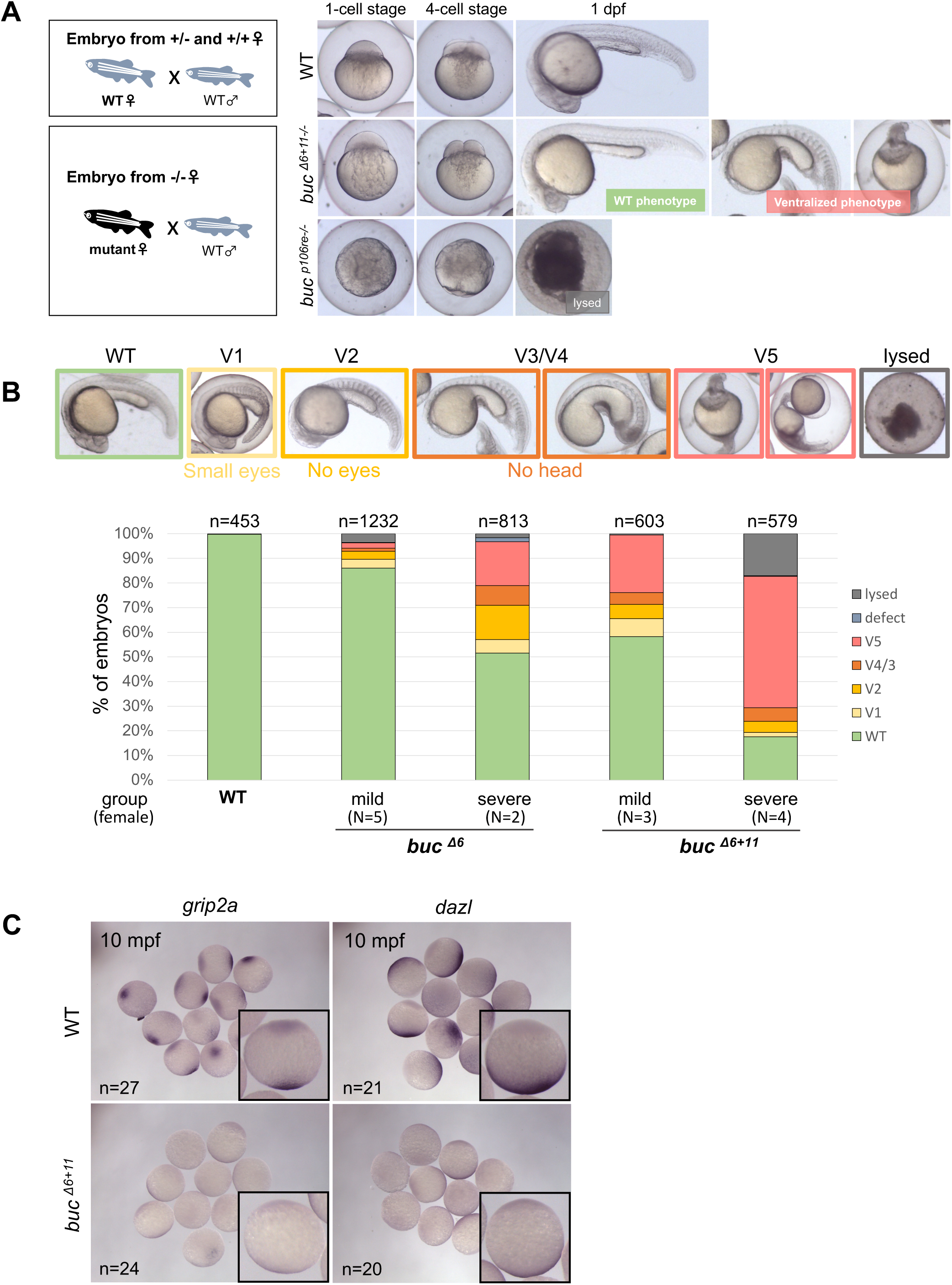
*buc* hypomorphic mutants have a normal A-V axis but severely ventralized embryos. **(A)** Embryos were obtained from wild-type (WT) and *buc* homozygous females. Also see Figure S2A-B. **(B)** Ventralized phenotypic distribution from *buc* CRISPR mutant females. Upper panel shows phenotypic classification of embryos at 1 dpf (V1 to V5: mild to severe phenotype). V1: Small eyes and blocky somites. V2: No eyes and blocky somites. V3 and V4: No head and blocky somite. V5: No yolk extension. Embryos (n) were obtained from multiple different females (N) of each group. WT embryos were obtained from 6 different crosses. Also see Figure S2C-D. **(C)** *In situ* hybridization of the vegetally-localized mRNA *grip2a* and *dazl* at 10 mpf. Embryos (n) were obtained from 3 different females; the right corner box shows a representative embryo.

mRNAs important for dorsal-ventral (D-V) axis formation localize in a Buc-dependent manner to the vegetal pole of the egg and early embryo^5–8^. In the *buc^p106re^* null mutant, vegetally-localized mRNAs are unlocalized in oocytes and early embryos^20^. Hence, we investigated the localization of vegetal mRNAs in embryos from the stronger *buc^Δ6+11^* hypomorphic mutant that produced a large fraction of ventralized embryos (Fig. 2B). We performed *in situ* hybridization for the vegetally-localized mRNAs, *glutamate receptor interacting protein 2a* (*grip2a*) and *DAZ-like* (*dazl*) in 10-minute post fertilization (mpf) embryos^7,24^. In embryos from *buc^Δ6+11^* mutants, both *grip2a* and *dazl* mRNA were localized at the vegetal pole but appeared reduced compared to wild-type (WT) (Fig. 2C). These results indicate that reduced vegetal pole localization of factors acting in D-V axis formation, such as *grip2a*, cause the ventralization phenotype of *buc^Δ6+11^* embryos.

### The Bb fails to form but Buc and *dazl* localize vegetally in later stage oocytes in *buc* hypomorphic mutants

Transcripts of *grip2a* and *dazl* first localize in a Buc-dependent manner to the Balbiani body (Bb) in stage I oocytes and then to the vegetal cortex at the end of stage I, as the Bb disassembles at the prospective vegetal pole (Fig. 3A). Hence, we examined stage I oocytes for the localization of Buc and *dazl* to the Bb. In WT stageⅠoocytes, Buc protein and mitochondria localized to the Bb, however, in both *buc^Δ6^* and *buc^Δ6+11^* stage I oocytes, we did not observe Buc or mitochondria aggregated into a mature Bb, although we found aberrant Buc aggregation in ∼10% of *buc^Δ6^* stage I oocytes (Fig. 3B). Next, we performed fluorescence *in situ* hybridization (FISH) for *dazl* mRNA in stage I oocytes. *dazl* mRNA was detected in the Bb in WT, but like Buc, we found that it was unlocalized in both *buc^Δ6^* and *buc^Δ6+11^* stage I oocytes (Fig. 3B). We next examined Buc and *dazl* localization in stage II oocytes, where in WT they reside at the vegetal cortex. Surprisingly, despite the Bb not forming in stage I *buc^Δ6^* and *buc^Δ6+11^* oocytes, Buc protein and *dazl* mRNA were localized at the vegetal cortex in *buc^Δ6^* and *buc^Δ6+11^* mutants (Fig. 3C, Fig. S3A-B). This was very unexpected because the dogma in the field is that the disassembling Bb is the conduit for Buc and multiple transcripts to localize to the oocyte vegetal cortex.

**Figure 3.**
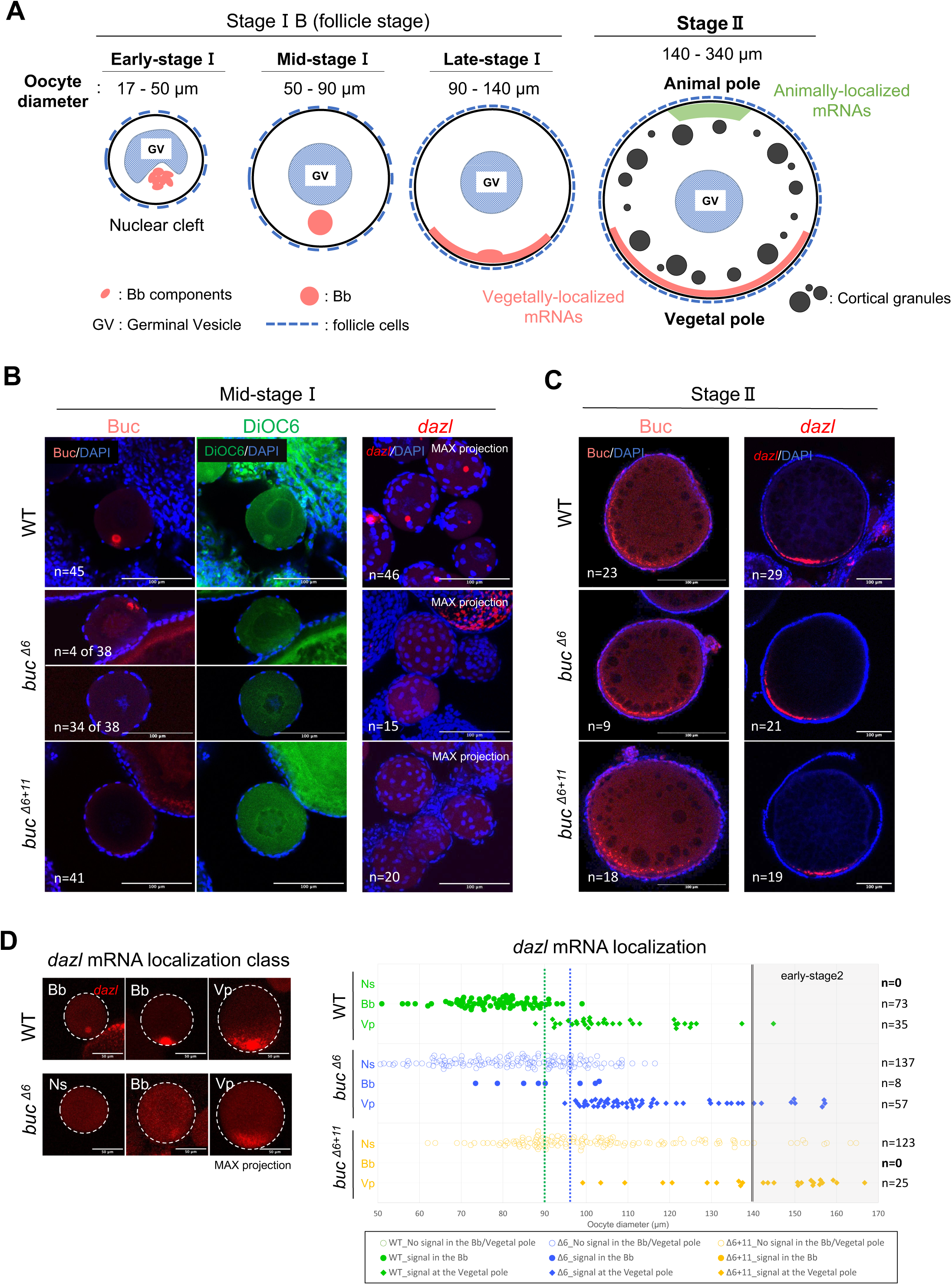
The Bb fails to form in stage I oocytes but Buc protein and *dazl* mRNA localized at the vegetal cortex of later stage *buc* hypomorphic mutant oocytes. **(A)** Bb formation and mRNA localization during zebrafish oogenesis. **Early-stage I (17-50 μm):** Bb component aggregates in the nuclear cleft. **Mid-stage I (50-90 μm):** matured round Bb is formed and grows correlated to the oocyte size. **Late-stage I (90-140 μm):** Bb starts disassembling and Bb components, such as Buc protein and *dazl* mRNA, become localized at the vegetal cortex. **Stage II (140-340 μm):** Other vegetal pole mRNAs and animal pole mRNAs localize at each pole. **(B)** Buc immunostaining (red, left) with DAPI (blue) and DiOC6 (green) staining, and fluorescence *in situ* hybridization (HCR) for *dazl* mRNA (red, right) with DAPI staining (blue) in mid-stage I oocytes. DAPI visualizes the nuclei of follicle cells surrounding oocytes. DiOC6 is a membrane dye and Bb marker because the Bb is a mitochondrial-rich structure. Buc staining was performed on oocytes from N=6 (WT), N=4 (*buc^Δ6^*), N=4 (*buc^Δ6+11^*) females and *dazl* staining was performed on oocytes from N=4 (WT), N=3 (*buc^Δ6^*), N=4 (*buc^Δ6+11^*) females. Oocytes (n) were imaged using a confocal microscope. Images of *dazl* mRNA staining were generated as max intensity projections. Scale bar = 100 μm. **(C)** Buc immunostaining (red, left) and HCR for *dazl* mRNA (red, right) in stage II oocytes stained with DAPI (blue). Buc staining was performed on oocytes from N=6 (WT), N=4 (*buc^Δ6^*), N=4 (*buc^Δ6+11^*) females and *dazl* staining was performed on oocytes from N=4 (WT), N=3 (*buc^Δ6^*), N=4 (*buc^Δ6+11^*) females, and oocytes (n) were imaged using a confocal microscope. Scale bar = 100 μm. Also see Figure S3A-B (late-stage I, early-stage II). **(D)** *dazl* mRNA localization in mid-stage I to early-stage II oocytes. z-stack images of oocytes were acquired to determine the oocyte diameter (X-axis) and *dazl* mRNA localization (Ns: No signal in the Bb/at the Vegetal pole, Bb: signal in the Bb, Vp: signal at the Vegetal pole). Oocytes (n) were obtained from 3 different females for quantification. White dotted-line outlines each oocyte. Green and blue vertical dotted line in the dot plot indicates the oocyte size at which *dazl* mRNA starts spreading at the vegetal cortex in WT and *buc*^Δ6^, respectively. Black double line (140 μm) indicates the oocyte size at stage II. Scale bar = 50 μm.

We then quantified *dazl* mRNA localization in mid-stageⅠto early-stage II oocytes (50-170 μm). In WT, *dazl* mRNA localized to the Bb in oocytes of diameter under 90 μm and then along the vegetal cortex in larger oocytes as the Bb disassembles (Fig. 3D). On the other hand, in *buc^Δ6^* and *buc^Δ6+11^* mutant, we rarely found a Bb-like signal (n=8 out of 194 in *buc^Δ6^* and n=0 in *buc^Δ6+11^*) but in *buc^Δ6^* stage I oocytes > 95 μm in diameter and *buc^Δ6+11^* stage II oocytes >140 μm in diameter, we found strong *dazl* mRNA signals along the vegetal cortex (Fig. 3D, Fig. S3A-B). These results suggest that a Bb-independent mechanism polarizes the oocyte by late-stage I of oogenesis.

### *dazl* is reduced vegetally and *cyclinB1* is expanded animally in *buc^Δ6+11^* but is normal in *buc^Δ6^*

In stage II oocytes (140-340 μm in diameter), animally-localized mRNAs begin to localize at the animal pole^25^ (Fig. 3A). In *buc^p106re^*null mutants, animal pole mRNAs expand their localization domain circumferentially around the oocyte cortex^20,26^. We examined animal-vegetal mRNA localization in *buc* hypomorphic mutant oocytes by FISH double-staining for animal pole mRNA *cyclin B1* ^25^ and vegetal pole mRNA *dazl* in stage II oocytes (Fig. 4A, Fig. S3B). In WT and *buc^Δ6^* mutants, we found that *cyclin B1* mRNA was normally localized at the animal pole and *dazl* mRNA was at the cortex encompassing the vegetal half of stage II oocytes (Fig. 4A). Interestingly, in *buc^Δ6+11^* stage II oocytes, *cyclin B1* mRNA localization was expanded animally, while the *dazl* mRNA domain appeared reduced compared to WT and *buc^Δ6^* (Fig. 4A).

**Figure 4.**
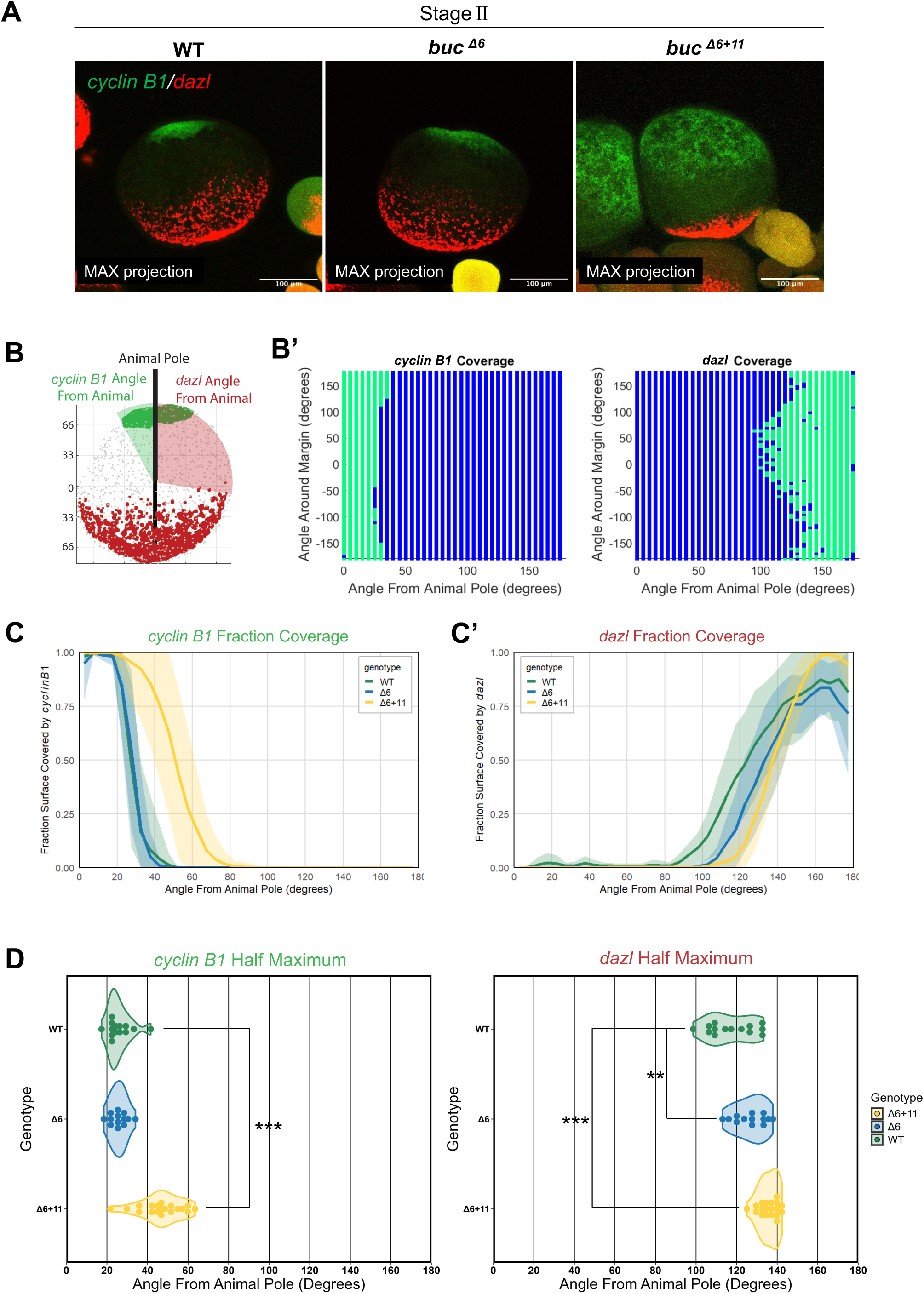
The *dazl* domain is reduced vegetally and *cyclin B1* domain is expanded animally in *buc^Δ6+11^* whereas *buc^Δ6^* appears normal. **(A)** HCR for *cyclin* B1 (green) and *dazl* (red) mRNAs in stage II oocytes (140-340 μm). *cyclinB1* localizes animally and *dazl* localizes vegetally. Images were generated as max intensity projections. Scale bar =100 μm. **(B)** The oocyte bounds (gray), *cyclin B1* domain (green), and *dazl* domain (red) were identified in 3 dimensions using an image processing pipeline in MATLAB. The animal pole was identified as the center of the *cyclin B1* domain. The oocyte is rotated so that the animal pole is upward (0 degrees) and the vegetal pole is downward (180 degrees). The black line is the A-V axis. The angles of the *cyclin B1* and *dazl* boundaries from the animal pole were measured. **(B’)** shows representative data from one oocyte. **(C)** The mean fraction of oocyte surface covered by *cyclin B1* **(C)** or *dazl* **(C’)** was calculated every 5 degrees from the animal to the vegetal pole for WT (n=15), *buc^Δ6^* (n=14), and *buc^Δ6+11^* (n=17). The ribbon is one standard deviation. **(D)** The *cyclin B1* and *dazl* fraction covered curves for individual oocytes were fitted with a sigmoidal curve to calculate the degree of the half-maximum of coverage. Each dot shows the individual oocyte half-maximum. Dunnett’s Test with WT as the reference was used to determine significance. ***= P < 10^-6, **= P < 10^-2.

To quantify the *cyclin B1* and *dazl* mRNA localization domains in *buc* hypomorphic mutant oocytes, we developed a quantitative analysis pipeline in MATLAB. We identified the oocyte surface, *cyclin B1* domain and *dazl* domain from acquired z-stack images and rotated the oocyte along the A-V axis (the animal pole: 0 degrees is defined as the center of the *cyclin B1* domain) and measured the *cyclin B1* and *dazl* boundary angles from the animal pole (Fig. 4B-B’). We calculated the mean fraction of oocyte surface covered by *cyclin B1* (Fig. 4C) and *dazl* (Fig. 4C’), and the degree of the half-maximum of coverage of individual oocytes (Fig. 4D). We found that the *cyclin B1* domain was significantly expanded in *buc*^Δ6+11^ mutant oocytes and the *dazl* domain was reduced in both *buc^Δ6^* and *buc^Δ6+11^*, but to a greater extent in *buc^Δ6+11^*.

### Mutant Buc protein can form small aggregates in early-stageⅠin *buc*^Δ6^ oocytes but fails to form a mature Bb

In early-stage I oocytes (17-50 μm in diameter), components of the Bb form multiple small aggregates around the centrosome and accumulate in a nuclear cleft prior to mature Bb formation^27^ (Fig. 3A). These premature Bb aggregates are in a liquid-like state and transitioning to the mature solid-like Bb observed in mid-stage I oocytes^28^. We investigated if these aggregates were formed in the *buc* hypomorphic mutants. In early-stage I oocytes (17-30 μm in diameter), Buc protein and *dazl* mRNA localized in the nuclear cleft in WT and *buc^Δ6^* mutants, while they were not detected in the nuclear cleft of *buc^p106re^*or *buc^Δ6+11^* mutants (Fig. 5A). In early to mid-stage I oocytes (35-50 μm in diameter), however, we did not observe a maturing Bb in *buc*^Δ6^ mutants, although dispersed Buc signal associated with DiOC6 next to the germinal vesicle (Fig. 5B). These results indicate that mutant Buc^Δ6^ protein can form small aggregates in early-stage I oocytes but fails to aggregate into a mature Bb. While the Buc^Δ6+11^ protein is predicted to be the same 10-amino acid N-terminal truncation as Buc^Δ6^, it fails to aggregate throughout stage I of oogenesis.

**Figure 5.**
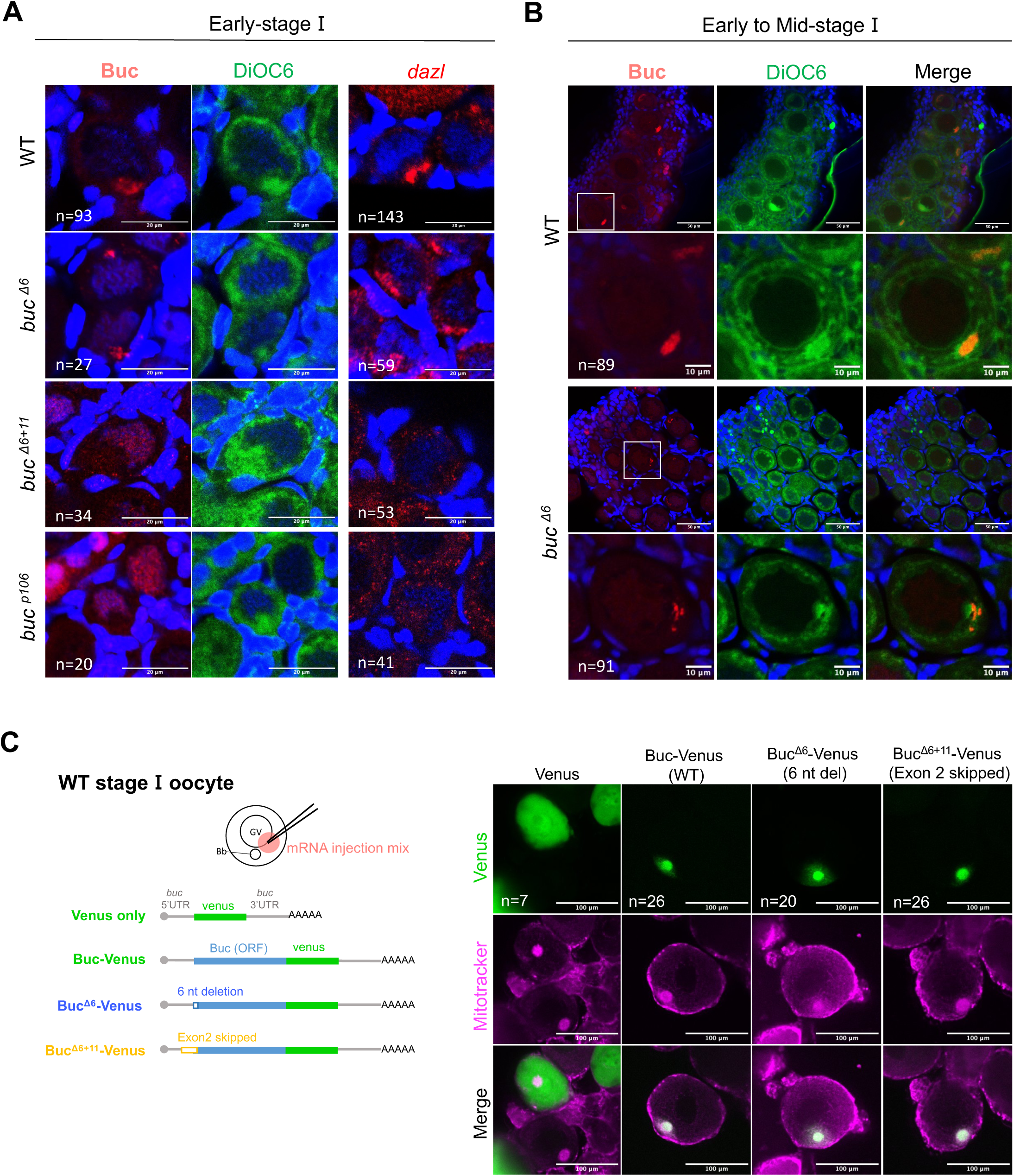
Mutant Buc protein can form pre-Bb aggregates in early-stage I *buc^Δ6^* oocytes but fails to form a mature Bb. **(A)** Buc immunostaining (red, left) in early-stage I oocyte stained with DAPI (blue) and DiOC_6_ (green), and HCR for *dazl* mRNA (red, right) in early-stage I oocyte stained with DAPI (blue) in juvenile gonad. N=5 WT, N=3 *buc^Δ6^*, N=3 *buc^Δ6+11^*, N=3 *buc^p106re^* were stained for Buc and N=4 WT, N=3 *buc^Δ6^*, N=3 *buc^Δ6+11^*, N=2 *buc^p106re^* were stained for *dazl*, and oocytes (n) were imaged using a confocal microscope. Scale bar = 20 μm. **(B)** Buc immunostaining (red) in early to mid-stage I oocytes stained with DAPI (blue) and DiOC_6_ (green) in a juvenile gonad. N=5 WT, N=3 *buc^Δ6^*, N=3 *buc^Δ6+11^*, N=3 *buc^p106re^* were stained for Buc and N=4 WT, N=3 *buc^Δ6^*, N=3 *buc^Δ6+11^*, N=2 *buc^p106re^* were stained for *dazl*, and oocytes (n) were imaged using a confocal microscope. Images below are a representative oocyte indicated in the white box in the upper panel. Scale bar =50 μm and =10 μm in cropped image. **(C)** Buc-venus and mutant Buc-venus localize to the Bb in WT stage I oocytes. Injected *buc^Δ6^-venus* mRNA has a 6 nt deletion and *buc^Δ6+11^-venus* mRNA lacks exon 2.

We next investigated whether the N-terminally truncated Buc^Δ6^ and Buc^Δ6+11^ hypomorphic proteins could aggregate into the Bb with WT Buc. To test this, we injected mRNA encoding Buc^Δ6^ or Buc^Δ6+11^ fused to Venus into WT stage I oocytes. We found that both N-terminally truncated Buc proteins could localize to the Bb in the presence of endogenous WT Buc protein (Fig. 5C). Thus, the levels of the Buc truncated protein may be insufficient to form a Bb on its own and/or the N-terminal 10 amino acids may be essential for its self-aggregation into the Bb, which is obviated when WT Buc is present.

### Buc mutant protein is reduced and can form a Bb when overexpressed but is often aberrant

Due to the mutated start codon in the hypomorphic *buc* mutants, translation is predicted to initiate 10 amino acids downstream at the next in-frame ATG codon, which displays a weak Kozak sequence^29–31^ (Fig. S1B). Thus, we hypothesized that the cause of the *buc* hypomorphic phenotype is due to (1) a reduced amount of Buc protein and/or (2) partial-loss of function due to the N-terminal Buc truncation. To address these hypotheses, we first tested the expression levels of endogenous *buc* mRNA in the *buc* lines: *buc^p106re−/−^*, *buc^p106re+/−^*, *buc^Δ6−/−^*, and *buc^Δ6+11−/−^*. We performed quantitative reverse transcription PCR for *buc* mRNA using total RNA extracted from stage IV oocytes. We found that the mutant *buc* mRNA is stable in *buc^Δ6^*, while it is significantly reduced in *buc^Δ6+11^* (Fig. 6A). The *buc^Δ6+11^* transcript skips exon 2, removing 51 nucleotides from the 5’ UTR, which may cause instability of the *buc^Δ6+11^* transcript.

**Figure 6.**
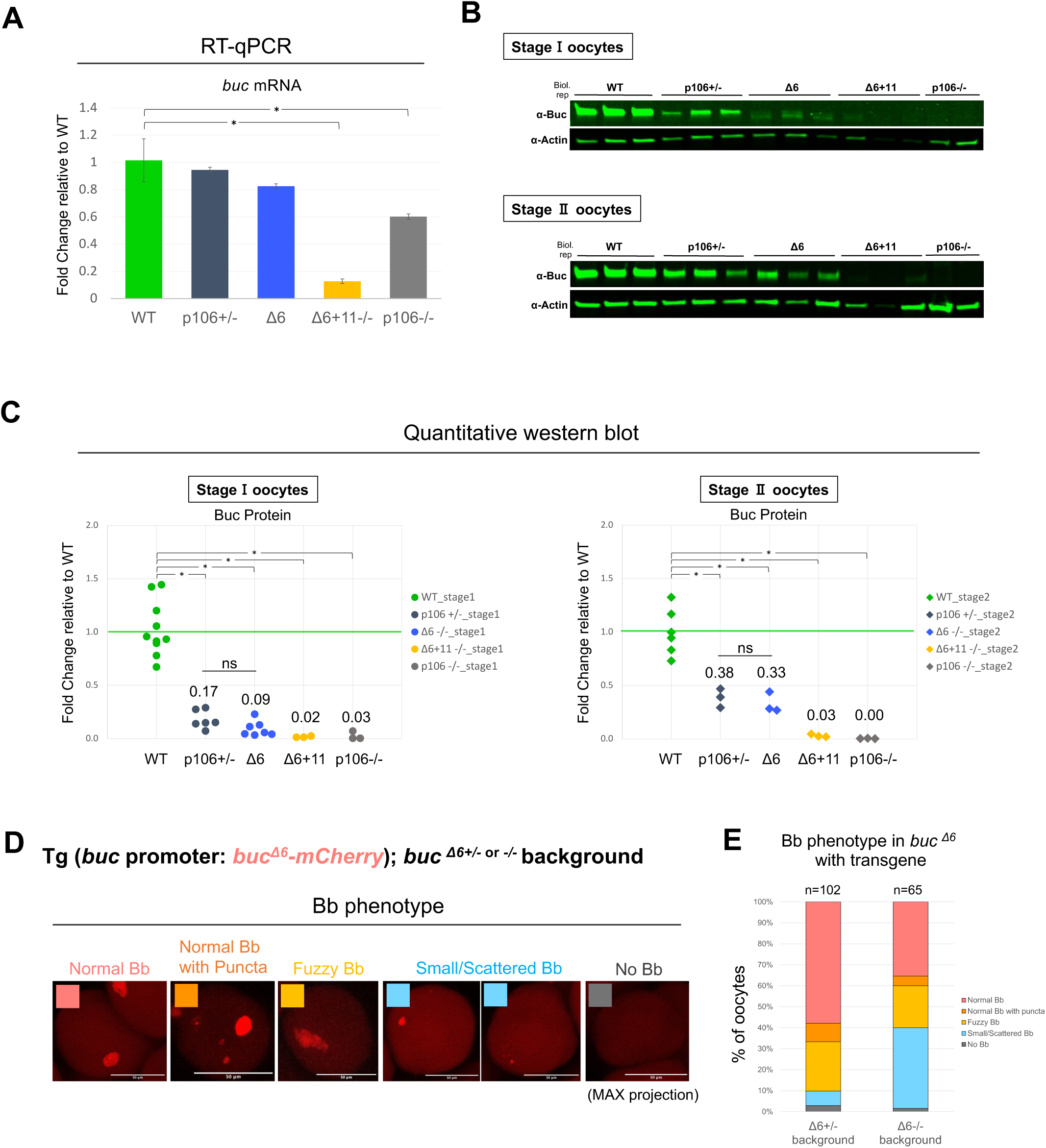
Buc mutant protein amount is reduced and can form the Bb when overexpressed in a transgene but the Bb is generally abnormal. **(A)** Quantitative reverse transcription PCR for *buc* mRNA. Total RNA was extracted from stage Ⅳ oocytes. Relative fold gene expression to WT was calculated by the delta-delta Ct method. *ef1a* was used as a reference gene. qPCR was performed for 3 biological replicates and the error bar indicates standard deviation. Significant difference between WT is shown as * (Dunnett’s test *α < 0.05). **(B)** Representative blots used for quantitative western blot analysis. Stage I and stage II oocytes were isolated from adult females and filtered by size using cell strainers (stage I: < 100 μm, stage II: 100-300 μm). anti-Actin was used as a positive control. See Figure S4A for total protein staining. **(C)** Quantitative western blot for Buc protein in stage I and stage II oocyte samples. Each dot indicates data points from different females. Buc signal was normalized by total protein staining and fold change relative to the average of WT females (set as 1) was calculated. Values above the plot are the average fold change of biological replicates. Significant difference is shown as * (p < 0.005, Welch’s t-test after Bonferroni correction). **(D)** Bb phenotype of Tg (*buc* promoter*: buc^Δ6^-mCherry*) in *buc ^Δ6^*. Bb phenotype in transgenic oocyte (*buc ^Δ6+/−^* or *buc ^Δ6−/−^* background) were classified into 5 classes (Normal Bb/ Normal Bb with puncta/ Fuzzy Bb/ Small and Scattered Bb/ No Bb). Images were generated by max intensity projections. Scale bar = 50 μm. **(E)** Quantification of the Bb phenotype in *buc^Δ6^* with Tg (*buc* promoter*: buc^Δ6^-mCherry*). Oocytes (n) were obtained from 5 different females for each genotype (*buc^Δ6+/−^* or *buc^Δ6−/−^* background).

Next, we performed quantitative fluorescent western blots to quantify the amount of Buc protein in stage I oocytes and stage II oocytes. Stage I and stage II oocytes were isolated from adult ovaries and sorted by size. In both stage I and stage II oocytes, Buc protein levels were significantly reduced in the *buc* hypomorphic mutant lines (Fig. 6B-C). Interestingly, the Buc protein level in stage I and II oocytes of *buc^Δ6^* was similar to that of *buc^p106re^* heterozygotes (Fig. 6C). Since the Bb forms in *buc^p106re^*heterozygotes but does not in *buc^Δ6^* oocytes, despite similar Buc levels, it suggests that the loss of the first 10 amino acids of the Buc^Δ6^ protein are important for Bb formation.

To test if the reduced levels of Buc^Δ6^ protein also contribute to the loss of Bb formation, we established a transgenic line to express more Buc^Δ6^ protein to determine if it could rescue the *buc^Δ6^* phenotype. We generated a transgenic line expressing mutant *buc*^Δ6^ fused to *mCherry* under the *buc* promoter in a *buc^Δ6^* background (Tg (*buc* promoter*: buc^Δ6^-mCherry*); *buc^Δ6^*, Fig. S4A). We then examined *buc^Δ6^* homozygous stage I oocytes with the transgene and, interestingly, found that about 40% of the oocytes displayed a normal or near normal Bb (Fig. 6D-E). This result suggests that the N-terminally truncated mutant Buc protein itself can form the Bb when overexpressed in a transgene. However, we also observed a large fraction of stage I oocytes with an abnormal Bb/Buc aggregates in transgenic females in a *buc^Δ6^* homozygous and heterozygous background (Fig. 6D-E, also see Fig. S4B for Bb size). Observing abnormal Bb or Buc aggregates also in the heterozygous background suggests that overexpression of mutant Buc protein can disrupt Buc self-aggregation.

In conclusion, our data show that *buc* hypomorphic mutants fail to form a mature Bb due to a reduced amount of Buc protein and to the loss of the first 10 amino acids of Buc. Normal A-V polarity is established in *buc* hypomorphic mutants lacking Bb formation, demonstrating that the Bb is not required for A-V polarity formation, but Buc protein itself plays a key role in establishing oocyte polarity.

## DISCUSSION

From studies in Xenopus and zebrafish, the Balbiani body (Bb) is believed to be an mRNA transport organizer key to localizing vegetal mRNAs important for embryogenesis to the oocyte cortex. Vegetally-destined transcripts reside within the Bb and at the end of stage I of oogenesis, the Bb disassembles at the prospective vegetal cortex, where the vegetal transcripts become docked. Analysis of *buc* null mutant oocytes supports the function of the Bb in vegetal mRNA localization and establishing animal-vegetal polarity during oogenesis. In *buc* null mutant oocytes, the Bb fails to form and vegetal mRNAs fail to localize to the vegetal cortex, while animal pole mRNAs expand radially at the oocyte cortex^20,26^. Here, we established through analysis of *buc* hypomorphic mutants that the Bb is not required for vegetal mRNA localization or oocyte polarity formation, shifting this paradigm of the field.

### The Bb and Bucky ball function in animal-vegetal polarity establishment

In a *buc* null mutant, the Bb fails to form and mutant oocytes and eggs lack animal-vegetal polarity^19,20^. On the other hand, our new *buc* hypomorphic mutant *buc^Δ6^* also fails to form the Bb (Fig. 3B and 3D) but remarkably displays normal animal-vegetal axis polarity (Fig. 2A and Fig. S2A-B). Moreover, although the Bb does not form, Buc protein and *dazl* mRNA, which was thought to localize vegetally via the Bb, are localized extensively vegetally in stage II oocytes (Fig. 3C, Fig. S3 and Fig. 4). These unexpected results show that the vegetal pole localization mechanisms and A-V polarity formation are established independently of Bb formation. Consistent with our results, previous studies found that a transgenic Buc protein that is unable to interact with Tdrd6a fails to form the Bb in a *buc* null mutant background, while the embryos still exhibit a normal A-V axis^32^. Thus Buc protein itself plays a key role to establish A-V polarity, but the Bb does not.

In Xenopus, a post-Bb stage vegetal localization pathway called the “late pathway” localizes mRNAs in a microtubule-dependent manner to the vegetal pole^17,33^. In Xenopus, some Bb-localized vegetal mRNAs injected into later-stage oocytes can localize to the vegetal cortex via the late pathway, indicating that these two pathways utilize overlapping mechanisms^34–38^. In zebrafish, *brul* and *mago-nashi* mRNAs normally do not localize to the Bb in stage I oocytes but do localize to the vegetal pole in post-Bb oocyte stages, stage II and III, respectively^20,39^. In *buc* null mutants, *brul* and *mago-nashi* fail to be localized vegetally, showing that Buc acts directly or indirectly in a later localization pathway(s). However, *dazl* mRNA and Buc protein localize in late stage I oocytes at the vegetal cortex in *buc^Δ6^* oocytes (95 μm diameter) (Fig 3D and Fig. S3A), which is earlier than this later localization pathway(s) (stage II, III, >140 μm diameter oocytes), suggesting that Buc and *dazl* are not untilizing this later pathway. Further research is needed to understand how the Bb components are localized at the vegetal cortex in zebrafish late-stage I oocytes.

### Bucky ball function in embryonic body axis formation

Embryos from *buc* hypomorphic mutant females display a ventralized phenotype, with *buc^Δ6+11^* exhibiting a more severe phenotype than *buc^Δ6^* embryos, which are mildly affected (Fig. 2B and Fig. S2C-D). Both mutants express the same predicted 10-amino acid truncated protein, but the Buc level is much more reduced in *buc*^Δ6+11^ than in *buc*^Δ6^ oocytes (Fig. 6A-C). These results suggest that the Buc protein amount may affect the ventralized phenotype severity. Gene transcripts acting in dorsal-ventral (D-V) axis formation, *syntabulin, grip2a,* and *huluwa,* localize to the Bb in stage I oocytes and are anchored at the vegetal cortex in later stage oocytes and eggs^6–8^. In embryos from *buc^Δ6+11^* mutant females, *grip2a* mRNA is still localized at the vegetal pole but is reduced (Fig. 2C). These results show that Buc protein is important for localizing body-axis-formation related mRNAs at the vegetal cortex and proper localization of these components is important for embryonic body axis establishment events. Also, we note that embryos from *buc^Δ6^* mutant females show a ventralized phenotype even though a similar level of Buc protein is detected in *buc^p106re^*heterozygous and *buc^Δ6^* homozygous stage 1 and stage 2 oocytes (Fig. 6C). These results suggest the importance of the first 10 amino acids of Buc in its function in body axis related mRNA localization.

### First 10 amino acids of Buc and protein levels in Bb formation

The Bb is a stable amyloid-like solid structure in Xenopus and zebrafish^21,28^. Buc and the *Xenopus* Buc ortholog, Xvelo1, are highly disordered proteins, containing a prion-like domain at its N-terminal region. In vitro self-assembly of Xvelo1 depends on its prion-like domain^21^. In a recent study, it is reported that Buc phase-separates into dynamic liquid-like granules in the nuclear-cleft of early-stage I oocytes in zebrafish and then condenses into a solid-like mature Bb^28^. In *buc^Δ6^* early-stage I oocytes, Buc protein and *dazl* mRNA are detected in the nuclear cleft and Buc protein formed aggregates but did not condense into a mature Bb in mid-stage I oocytes (Fig. 5A-B). We confirmed that injected mutant Buc protein can associate with the Bb in a WT background (Fig. 5C). Additionally, we analyzed Buc protein sequence with the prion-like domain prediction algorithm, PLAAC^40^, and confirmed the deletion of the first 10 amino acids leaves intact the prion-like domain with similarity to that of Xvelo (Fig. S5A).

Protein concentration is one key factor that can contribute to phase transition, so the reduced amount of Buc protein in *buc^Δ6^* may contribute to failure of Bb formation (Fig. 6B-C). When we overexpressed Buc^Δ6^ protein via a transgene in a *buc*^Δ6^ mutant background, a nearly normal Bb could form, but also abnormal Bb/Buc aggregates were observed (Fig. 6D-E). These results indicate that the Buc^Δ6^ protein itself has the ability to aggregate into a Bb when overexpressed. However, Buc^Δ6^ overexpression did not fully rescue Bb formation, indicating that Buc^Δ6^ function is compromised. Furthermore, interestingly, we observed abnormal Bb and Buc aggregates in both a *buc^Δ6^* homozygous and heterozygous background (Fig. 6D-E, Fig. S4C-C’), suggesting that overexpression of Buc^Δ6^ protein disrupts wild-type Buc aggregation.

Multiple of our results indicate that truncation of the first 10 amino acids of Buc^Δ6^ compromises its function. As mentioned, a Buc^Δ6^ transgene fails to fully rescue the *buc*^Δ6^ mutant and induces a phenotype in *buc*^Δ6^ heterozygous oocytes, unlike a WT Buc transgene in *buc* null mutants^23^. Second, we detected a similar level of Buc protein in *buc^Δ6^* stage I and stage II oocytes as in *buc^p106re^*heterozygotes (Fig. 6C), however, *buc*^Δ6^ mutants lack the Bb, whereas *buc^p106re^*heterozygotes display a normal Bb. Buc protein sequence alignment among species shows no conservation among the first 10 amino acids, but this region is a predicted disordered region in PONDR^41^ (Fig. S5B). We hypothesize that the first 10 amino acids of Buc are important for (1) self-aggregation, (2) protein stability, (3) interaction with a binding partner, and/or (4) for post-translational modification. Future studies are required to determine the contribution of these potential Buc^Δ6^ attributes.

### What is the function of the Bb condensate in oocyte polarity formation?

It is postulated that the Bb is a biomolecular condensate that directs its localized components to the vegetal cortex and establishes oocyte polarity in Xenopus and zebrafish. However, unexpectedly, Bb-localized components localized at the vegetal cortex without Bb formation in *buc* hypomorphic mutants in our study. As discussed above, Xenopus Bb-localized mRNA injected into the post-Bb stage oocyte can be localized at the vegetal cortex, also indicating that their association with the Bb is not essential for their vegetal localization. The timing of *dazl* mRNA and Buc protein localization at the vegetal cortex in *buc^Δ6^* mutant (late-stage I, 95 μm diameter) (Fig 3D and Fig. S3A) corresponds to the timing of Bb disassembly in WT, suggesting that the vegetal cortex localization of the Bb components may be achieved via a Bb-independent indirect translocation mechanism. Also, it is reported that *syntabulin* mRNA becomes unlocalized in *buc^p106re^* but a similar level of mRNA as in WT oocytes is detected by qRT-PCR^6^, and *vasa, dazl, brul* and *nago-nashi* mRNA are not degraded in *buc^p106re^* mutant oocytes as well^20,26^. These results indicate that the Bb/Buc does not affect the stability of transcripts normally vegetally-localized in the oocytes. In recent years, a number of studies have addressed the function of biomolecular condensates as a specialized compartment for RNA biogenesis, but there is also emerging counterevidence suggesting that some condensates arise as by-products during RNA-protein assembly and serve no function^42^. Our study is another example that suggests the formation of a condensate itself, in our case, the Bb, is not important for the cellular function, at least mRNA translocation or polarity formation.

In conclusion, the *buc* hypomorphic mutants we established in this study demonstrate that the Bb is not required for oocyte A-V polarity formation, which is a paradigm shift for the field. It is clear, however, that Buc itself plays a key role in oocyte polarity establishment. While *buc^Δ6^* early stage I oocytes at a pre-Bb stage display Buc^Δ6^ and *dazl* asymmetry, possibly reflecting polarity establishment by Buc, *buc ^Δ6+11^* early stage I oocytes do not show such asymmetry. However, both mutants exhibit clear polarity at the end of stage I of oogenesis, when Buc and *dazl* both localize to the vegetal pole. These results raise questions about whether a factor(s) upstream of Buc in early stage I oocytes, functions with Buc to polarize the oocyte, and/or if a window of competence to establish polarity exists until at least the end of stage I of oogenesis when asymmetric Buc^Δ6+11^ protein is first apparent. Many questions remain, which will require further studies.

## ACKNOWLEDGMENTS

We would like to thank the Cell & Developmental Biology Microscopy Core, especially Andrea Stout and AJ, for helpful support and past and present members of the Mullins lab. This work was supported by a grant from the National Institutes of Health (R35-GM131908).

## AUTHOR CONTRIBUTIONS

M.K. and M.C.M. conceptualized and wrote the manuscript. M.K. conducted the experiments. J.Z. developed MATLA pipelines for the oocyte domain size quantification. M.K. and J.Z. performed the analysis and generated the figures.

## METHODS

### EXPERIMENTAL MODEL AND SUBJECT DETAILS

#### Zebrafish Lines

Tü (Tübingen) strain was used as the background of the *buc* CRISPR mutants and the transgenic line. Fish lines used in this research were: Tü (WT), *buc^p106re^* ^19^, *buc^p6del^*, *buc^p6del11in^*, Tg (*buc*: *buc ^p6del^*-*mCherry*-*buc* 3’UTR); *buc^p6del^ or buc^p6del^/+*. All zebrafish strains were maintained at 28 °C with an 11/13-hour dark/light cycle. Embryos were kept and raised at 28 °C.

#### Ethics statement

The animal study was reviewed and approved by the University of Pennsylvania Perelman School of Medicine Institutional Animal Care and Use Committee.

### METHOD DETAILS

#### CRISPR/Cas9 mutagenesis

*buc* CRISPR mutants were generated using the Alt-R™ CRISPR-Cas9 System (IDT). crRNA (IDT) was designed in exon2, around start codon using CHOPCHOP^43^. crRNA was incubated with tracrRNA (IDT) to form a functional gRNA duplex in Nuclease-free duplex buffer (IDT). Then gRNA duplex was mixed with Cas9 protein (PNA Bio, Cat# CP02-250) to form an RNP complex. 1-3 nl of 2.5 μM RNP complex with 0.05 % phenol red/100 mM KCl was injected to 1-cell stage embryos.

To identify mutation carrier fish, we performed the HRM (High Resolution Melting analysis) using MeltDoctor HRM Master Mix (Applied Bio-systems, Cat# 4415440). Primer set used in HRM are listed below.

***buc* DNA target sequence used for crRNA**
5’ GTCTCCTCGTAGGAAATGGA 3’

***buc* HRM Fwd.**
5’ ATCCTGTGCTGATTTTCTTCGT 3’

***buc* HRM Rev.**
5’ AGGTGAATAGTTAATCTGAACCGT 3’

#### Sequencing

To identify the mutation in *buc* CRISPR mutants, we extracted gDNA from an F1 embryo or a piece of tail fin^44^. Tail fins were clipped from adult fish anaesthetized with 0.016% SYNCAIN (Syndel USA, Cat# NC0872873) in fish system water. We performed genomic PCR and PCR product were purified with QIAquick Gel Extraction Kit (QIAGEN, Cat# 28706) for sequencing. Primers used in gPCR and sequencing are listed below.

***buc* N-term seq. Fwd.**
5’ CTGCTGTCGTCAAAACACTC 3’

***buc* N-term seq. Rev.**
5’ TGCCAAGTCCTTGTCATCC 3’

#### RT-PCR

Total RNA was extracted from unfertilized eggs (n=30) using TRIzol reagent (Invitrogen, Cat# 15596026) following the manufacturer’s protocol. Using SuperScript III First-Strand Synthesis System (Invitrogen, Cat# 18080051), cDNA was synthesized from 500 μg of total RNA with oligo dT primer. RT-PCR and sequencing of PCR products were performed using the primers below.

***buc* exon1 Fwd.**
5’ TGGATCTCTGGAAACAGACG 3’

***buc* exon3 Rev.**
5’ CTTGTGTGGTTTACTGGGTG 3’

#### Genotyping

We genotyped *buc* CRISPR mutants using KASP^TM^ genotyping technology (LGC Bioresearch Technologies). Flanking sequences used for designing KASP assay primers are listed below.

**[WT/mutant]**

***\*buc^p6del^***
ACCGTGTACTGACATTGTTTGGTCTAATCTAAGTTTTGTACTTTATCTAGTATTGCGGTGTTTTGTATCCTGTGCTGATTTTCTTCGTCTCCTCGTAG[GAAATG/-]GAAGGTAA TCTTCAGAAACTTTTATTCTGCTAAAGCAGTTGAATACATTT

*\*buc^p6del11in^*
AGTATTGCGGTGTTTTGTATCCTGTGCTGATTTTCTTCGTCTCCTCGTAG**[GAAATG/ AAGGAGGAGAC]**GAAGGTAATCTTCAGAAACTTTTATTCTGCTAAAGCAGTTGAATA CATTT

For genotyping Tg (*buc*: *buc ^p6del^*-*mCherry*-*buc* 3’UTR); *buc^p6del^*, we performed gPCR using WT *buc* locus-specific primer to identify the background genotype and primer designed in the *mCherry* sequence to identify transgene carriers.

***buc* exon2 Fwd. for WT *buc* locus (p6del not amplified)**
5’ TTCGTCTCCTCGTAGGAAATG 3’
***buc* N-term seq. Rev.**
5’ TGCCAAGTCCTTGTCATCC 3’

**mCherry Fwd.**
5’ CACGAGTTCGAGATCGAGGG 3’
***buc* exon7 Rev.**
5’ GCAGCAAGGTGTACATCACTGTC 3’

*For genotyping *buc* hypomorphic alleles, we also designed *buc* hypomorphic alleles specific primers and performed gPCR with *buc* N-term seq Rev. primer.

***buc^p6del^*Fwd.**
5’CGTCTCCTCGTAGGAAGG 3’

***buc^p6del11in^ Fwd.***
5’ TCCTCGTAGAAGGAGGAGAC 3’

#### Buc antibody

We generated rabbit anti-Buc antibody against Buc C-terminal region (#609-624: RPRSEYNDYDETEFTY) (YenZym Antibodies, LLC.).

#### Whole-mount colorimetric *in situ* hybridization

Embryos were fixed in 4% PFA/PBS (diluted from 16% PFA, Electron Microscopy Science, Cat# 15710) orver night (O/N) at 4 °C and gradually dehydrated in MeOH (25%, 50%, 75% MeOH/PBS and 100% MeOH) and stored at −20 °C. Whole mount in situ hybridization was performed as described^45^ except that hybridization was at 65°C. After rehydration and PBST (PBS with 0.1 % Tween20) wash, embryos were hybridized with DIG-labeled RNA probes for *dazl* ^24^, *grip2a*^7^, and detected with anti-DIG-AP (Rosch, Cat# 11093274910). For colorimetric detection, embryos were incubated in BM-purple (Rosch, Cat# 11442074001) substrate solution, and color development was checked under the microscope. Images were acquired with Leica IC80 HD camera.

#### Whole-mount fluorescent *in situ* hybridization and immunochemistry

Ovaries were dissected from adult females, and stage I and stage II oocytes were isolated manually. To observe early-stage I oocytes, juvenile gonads were obtained from juvenile fish (Standard Length^46^ = 1.0-2.0 cm) and oocytes and Juvenile gonads were fixed in 4% PFA/PBS O/N at 4 °C. *dazl* and *cyclin B1* were detected using HCR™ Amplification Technology^47^ (Molecular Instruments) following the manufacturer’s protocol^48^ (Molecular Instruments). Immunochemistry was performed as described^49^. Samples were washed with 0.5% PBT (PBS with 0.5% TritonX-100) and blocked in 10 % FBS/PBT, then incubated with the primary antibody for Buc (1:250, This paper) O/N at 4 °C. Secondary antibody used was anti-rabbit IgG Alexa 594 (1:500, Invitrogen Cat# A-11037). Nuclei were stained with DAPI (1:1000, 300 μM stock, Invitrogen Cat# D3571) and mitochondria were stained with membrane dye DiOC6 (1:5000, Invitrogen Cat# D273).

#### Confocal microscopy

Images were acquired on Zeiss LSM 880 confocal microscope. Oocytes and juvenile gonads were mounted in VECTASHIELD Antifade Mounting Medium with DAPI (Vector Laboratories, Cat# NC1848444) as described^49^ for imaging. We made an imaging well on a rectangle #1.5 coverslip (Fisherbrand Cat# 12-544-CP) using Secure-Seal™ Spacer (0.12mm deep, Invitrogen Cat# S24737) and covered with a square #1.5 coverslip (Fisherbrand Cat# 12-541-AP). For domain size quantification in stage II oocytes, we used 3 stacks of Secure-Seal™ Spacer to avoid squashing the oocytes. Oocyte samples were mounted in BABB (1:2 of benzyl alcohol and benzyl benzoate)^50^ and obtained z-stack images with 25x/0.8 oil immersion lens (2.6 μm interval, 1024 x 1024 pixels). Live oocyte samples dissected from adult transgenic females were imaged in the 35 mm glass-bottom dish (MatTek Cat# P35G-1.5-10-C) within L-15 medium (Gibco Cat# 11415064). Also see Quantification and statistical analysis section below.

#### Quantitative RT-PCR

For *buc* mRNA quantification, total RNA was extracted from manually isolated stage IV oocytes (n=50 each). cDNAs were synthesized as described in the previous section. qPCR was performed on Applied Biosystems QuantStudio3 using PowerUp SYBR Green Master Mix (Applied Biosystems Cat# A25742). *ef1a*^51^ was used for the reference gene and the relative expression level to WT was calculated using the delta-delta Ct method (Also see Quantification and statistical analysis section). Primer sets used in qPCR are listed below.

***buc* qPCR Fwd.**
5’ TGCAGGCCTTGTATTGAGC 3’
***buc* qPCR Rev.**
5’ TCCAGTAACTTGCCTCCAAGAC 3’

***ef1a* qPCR Fwd.**
5’ TGATCTACAAATGCGGTGGA 3’
***ef1a* qPCR Rev.**
5’ CAATGGTGATACCACGCTCA 3’

#### Oocyte isolation for western blot

For quantitative western blot, we isolated stage I and stage II oocytes from 1 adult female using the 300 μm and 100 μm cell strainers (PulriSelect Cat# 43-50300-03 and MTC Bio Cat# MTC-C4100) based on previous works^49,52^. To avoid yolk protein contamination, we removed yolk-containing oocytes manually before enzyme treatment, then digested them within the collagenase cocktail (mix of 3 mg/ml Collagenase I, 3 mg/ml Collagenase II, and 1.6 mg/ml Hyaluronidase in L-15) for 15-20 min at RT. After separating oocytes by gentle tapping and pipetting with p200 tip, the oocyte sample was passed through the stacked cell strainers (mesh size 300 μm and 100 μm) attached to the 6-well plate (VWR, Cat# 10861-554) to collect stage I oocytes (< 100 μm). Then flipped the 100 μm cell strainer to collect stage II oocytes (100-300 μm). Samples were washed 2 times with L-15 and 2 times with PBS and stored at −80 °C.

#### Quantitative fluorescent western blot

Oocyte samples were suspended in NP-40 lysis buffer with protease inhibitor (Rosch Cat# 04693124001) and homogenized with a micro pestle on ice. Samples were denatured with Laemmli sample buffer (BioRad Cat# 1610737) at 95 °C for 5 min before loading on a 4–15% Mini-PROTEAN TGX Precast Gel (BioRad Cat# 4561086) and blotted on a low-fluorescence PVDF membrane (BioRad Cat# 1620261) using XCell II Blot Module (Invitrogen, EI9051). For western blot normalization, total protein was stained with Revert™ 700 Total Protein Stain (LI-COR Cat# 926-11010) and imaged on a LI-COR ODYSSEY M. Following total protein staining, the membrane was cut in half for detecting Buc and Actin as a positive control. Membrane was rinsed with TBST (TBS with 0.1 % Tween20) and incubated with blocking buffer (1% drymilk/TBST for Buc, 2.5% drymilk/TBST for Actin) at RT for 1 h and then incubated with the primary antibody (anti-Buc, this paper, 1:000, anti-Actin, Sigma-Aldrich, Cat# A2066, 1:5000 diluted with 2% BSA/TBST) at 4 °C O/N. After washing with TBST, membranes were incubated with secondary antibody (a-Rabbit-IgG (H+L), DyLight 800 4xPEG, Invitrogen Cat# SA5-35571, 1:25,000 diluted with 1% drymilk/TBST) at RT for 1 h and imaged on a LI-COR ODYSSEY M for quantification.

#### mRNA injection

Oocyte injection was performed as described^53^ and imaged with Yokogawa CSU-X1 Spinning Disk Confocal (Olympus IX81 inverted microscope). mRNAs were synthesized with mMESSAGE mMACHINE™ SP6 Transcription Kit (Invitrogen Cat# AM1340). mRNA (800 ng/μl) with 0.05 % phenol red/100 mM KCl was injected about the same size as the diameter of the oocyte nucleus using the Sutter Instrument XenoWorks Digital Microinjector.

#### Cloning for the transgenic line

Tg (*buc*: *buc ^p6del^*-*mCherry*-*buc* 3’UTR); *buc^p6del^*was established using the tol2-transposon system^54,55^. 1 nl of 25 ng/μl of pmini-tol2*-buc: buc^p6del^-mCherry-buc* 3’UTR plasmid construct and 25 ng/μl of tol2 transposase mRNA^56^ in 200 mM KCl was injected into the cytoplasm of 1-cell stage embryos.

**[pmini-tol2-*buc^p6del^*5’ half]**
pmini-tol2-*tnnt2a:GM2-GFP* (pDB782, gift from Dr. Darius Balciunas^57^) was used as the backbone. To clone *buc* promoter^23^ and *buc* mutant sequence, we extracted genomic DNA from *buc^p6del−/−^* embryos (n=60) and used it as a PCR template. Embryos were incubated with lysis buffer (100 μg/ml proteinase K (Thermo Scientific Cat#EO0491), 20 μg/ml RNase (Rosch Cat#11119915001), 0.5% SDS) at 37 °C O/N, then extracted 2 times with phenol/chlorophorm/isoamylalcohl (Sigma-Aldrich, Cat# P2069) and performed ethanol precipitation with NaCl and then NaOAc. *buc* promoter and *buc* mutant sequence until Exon6 was amplified using the primer set below and ligated with the linearized backbone digested with Swa1 (New England Biolabs Cat# R0604S) using In-fusion cloning HD kit (Takara Bio Cat# 639650). Library efficiency DH5a competent cell (Invitrogen, Cat# 18263012) was used for transformation.

**In-fusion *buc* prom Fwd.**
5’ TGCCCAGTTTAATTTGCCAGCTCTTACCTCCTTCCTG 3’
**In-fusion *buc* Exon6 Rev.**
5’ TCGCCAGATCTATTTCTCTGCCCCTCTGGCAGTAG 3’

**[pGEM-T easy *buc* 3’half with mCherry]**
To construct *buc Exon6-7-mCherry*-*buc* 3’UTR, we cloned *buc* Exon6-*buc* 3’UTR region to pGEM-T Easy (Promega, Cat#A1360) and integrated *mCherry* sequence using In-Fusion technique. pCS2+Buc (CDS)-*mCherry* with *buc* UTRs was used as a template to amplify the *mCherry* sequence. The primer sets used here are listed below. Subcloning or library efficiency DH5a competent cells (Invitrogen, Cat# 18265017, 18263012) were used for transformation.

***buc* Exon6 Fwd.**
5’ GGAGAACTCCATTGAGAAGGTC 3’
***buc* 3’UTR Rev.**
5’ AGGTCTGCTCGTTAAAGTGG 3’

**In-fusion pGEM-T easy *buc* 3’half vector linearization Fwd.**
5’ TAAACCATTATGCACTGCTGT 3’
**In-fusion pGEM-T easy *buc* 3’half vector linearization Rev.**
5’ GTATCTTGAGCCTCTTTTCTTCA 3’

**In-fusion mCherry Fwd.**
5’ AGAGGCTCAAGATACATGGTGAGCAAGGGCGAGG 3’
**In-fusion mCherry Rev.**
5’ GTGCATAATGGTTTACTTGTACAGCTCGTCCATGCC 3’

**[pmini-tol2-buc: buc^p6del^-mCherry-buc 3’UTR]**
To generate the final construct for transgenic line establishment, pmini-tol2-buc 5’ half (vector) and pGEM-T easy buc 3’half with mCherry (Insertion) were digested with Sal1 (New England Biolabs Cat# R0138S) and Bgl2 (New England Biolabs Cat# R0144S) and ligated at 16°C O/N (New England Biolabs Cat# M0202). Subcloning efficiency DH5a competent cell (Invitrogen, Cat# 18265017) was used for transformation.

#### Protein sequence analysis and alignments

Protein sequence alignments were performed using UniProt with the Clustal Omega program ^58,59^. Buc ortholog protein sequences used for alignment are *Danio rerio* (H0WFA5, BUCKY_DANRE), *Xenopus laevis* (Q7T226, Q7T226_XENLA), *Oryzias latipes* (A0A8G1GNG9, A0A8G1GNG9_ORYLA), *Salmo salar* (A0A1S3LYM3, A0A1S3LYM3_SALSA), Alligator sinensis (A0A1U7RV87, A0A1U7RV87_ALLSI), *Gallus gallus* (A0A8V0Z3J5, A0A8V0Z3J5_CHICK), *Ornithorhynchus anatinus* (A0A6I8N8D4, A0A6I8N8D4_ORNAN), *Sus scrof* (A0A8D0YR31, A0A8D0YR31_PIG). Prion-like domain prediction was performed using PLAAC^40^ and the intrinsic disorder of proteins was predicted with PONDR-VLXT^41^.

### QUANTIFICATION AND STATISTICAL ANALYSIS

#### Oocyte and Bb diameter and oocyte domain quantification

Quantification of the Oocyte and Bb diameter was calculated using Fiji^60^ following previous papers^49,61^. The focal plane with the largest oocyte area and the Bb area was used to measure the length of the two perpendicular lines along the oocyte/Bb diameter, and the average of the two lines was used as the approximate diameter of the oocyte/Bb.

For stage II oocyte (*dazl, cyclin B1*) domain quantification, we established the pipeline using MATLAB. We analyzed three-channel 3D oocyte stacks (1024×1024×Z; TIFF; .czi converted in Fiji). Voxel sizes were 0.3321×0.3321×2.6 µm. Channel 1 (C1) was used to segment the oocyte; channels 2 and 3 captured *cyclinB1* (C2) and *dazl* (C3). Volumes were denoised with 3D Gaussian filters (C1: σ≈24×24×3 px; C2: 24×24×3; C3: 14×14×3). Oocyte segmentation (C1) applied global binarization (level = median + sd, slice-wise dilation and hole-fill), retained the largest 3D component, and refined boundaries with a broad mask and per-slice distance-transform watershed. For each slice we kept the object whose centroid lay closest to the image center, then computed a slice-wise convex hull; masks were lightly eroded and combined with hull completions generated after axis permutation to correct coverslip-related gaps. The final oocyte mask (minRadBW_CH_Tot) was used to zero background in all channels.

Within this mask we thresholded C2 and C3 to obtain marker-positive voxels, mapped each hull voxel to an elevation angle (0 - 180° from the animal pole), and binned by 5° to compute per-angle surface coverage (*cyclinB1*elePercC2, *dazl* elePercC3) and volume fractions (H1volume_all, H1volumeC2_all). Coverage curves were summarized by logistic fits: C2 c3/[1+exp(−c1(x−c2))] and mirrored C3 c3/[1+exp(−c1(−x+c2))], with starts (−0.5, 30, 1) and (−0.2, 120, 1) and bounds c1∈[−1,−0.09],c2∈[5,170],c3∈[0,1]; the midpoint c2 defined C2_Bound/C3_Bound. Visualization used mean ± SD coverage curves and violin+dot plots of boundary angles, with WT, Δ6, and Δ6+11 plotted in fixed color order (green, blue, yellow). Dunnett’s Test with WT as the reference was used to determine significance. ***= P < 10^-6, **= P < 10^-2.

#### Relative quantification of *buc* using qPCR

The relative expression level to WT was calculated using the delta-delta Ct method. 3 biological replicates for each genotype were used for statistical analysis. Using Thermo Fisher Connect Platform Data Analysis App (Thermo Fisher), the Ct value for *buc* and *ef1a* (reference gene) was determined. The baseline and threshold were adjusted if needed. The Ct value for each sample was calculated as the average of 3 technical replicates (wells). ΔCt and ΔΔCt value was calculated on Microsoft Excel (Microsoft) using the formula below,

ΔCt = Ct (*buc*) - Ct (*ef1a*)

ΔΔCt = ΔCt (*buc*) - Average ΔCt of Control group (WT)

Then, the fold change relative to WT was calculated as 2^ (-ΔΔCt). The average of 2 technical replicates (experiments), except for WT and *buc*^p106re^, was used as the fold change of each sample. For statistical analysis, Dunnett’s test was performed (*α = 0.05). The error bar indicates standard deviation.

#### Quantification of the fluorescent western blot

The quantification of fluorescent western blot was performed using Image Studio software (LI-COR). Signal for the protein of interest (Buc) and total protein staining were measured using the “Draw Rectangle” function, and the “Median, segment All, Border width 2-pixel” was used as the background method. Biological replicates for WT (n=10), *buc^p106re+/−^* (n=6)*, buc^pdel6^* (n=7), *buc^pdel6ins11^* (n=3), *buc^p106re−/−^* (n=3) for stage I and WT (n=6) and n=3 for other genotypes for stage II oocyte samples were used for statistical analysis. Buc signals were normalized with the total protein staining following the Revert Total Protein Stain Normalization Protocol (icorbio.com/support/contents/applications/western-blots/revert-normalization-protocol.html). The fold change was calculated using the average of the WT group as 1 within each blot. The fold change of each biological replicate was calculated as the average of the technical replicates (at least 2 blots for each sample). p-values were calculated using two-sided Welch’s t-test with Bonferroni correction (*p < 0.005).

## SUPPLEMENTAL FIGURE LEGENDS

**Supplemental figure 1, Related to figure 1.**
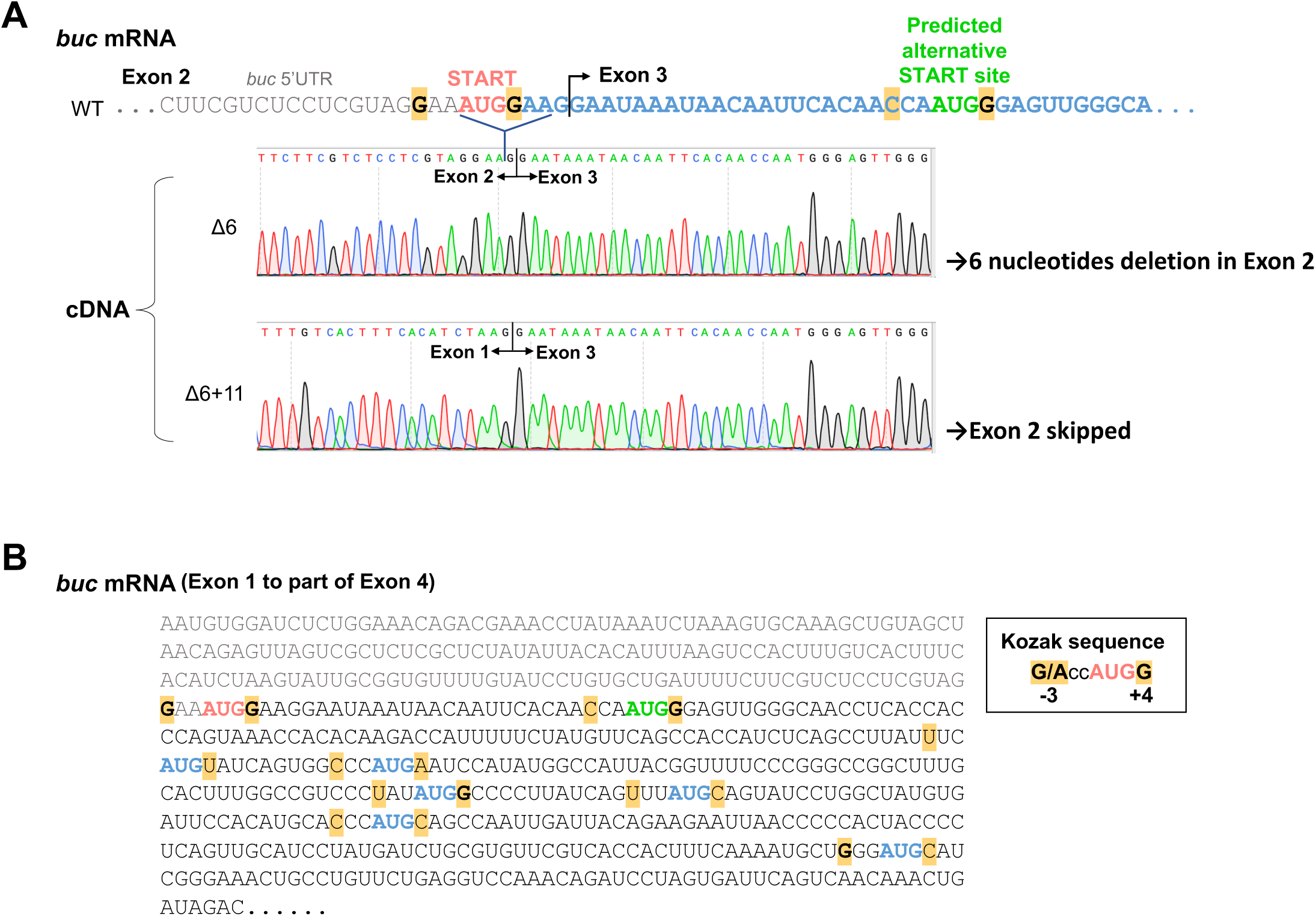
*buc* CRISPR mutant mRNA lacks start codon and predicted alternative start site is next AUG in exon 3 with weak Kozak sequence. **(A)** Sequencing results of maternal *buc* transcript from *buc* CRISPR mutant. **(B)** AUG in *buc* mRNA sequence. Red AUG indicates the start codon of *buc* in exon 2, and black bold letters indicate the Kozak sequence important for the translation efficiency. Position −3 and +4 are highlighted in yellow. Green AUG indicates predicted alternative start codon in exon 3 in *buc* CRISPR mutants. Blue AUG indicates other in-frame AUG in exon 3 and exon 4.

**Supplemental Figure 2, Related to figure 2.**
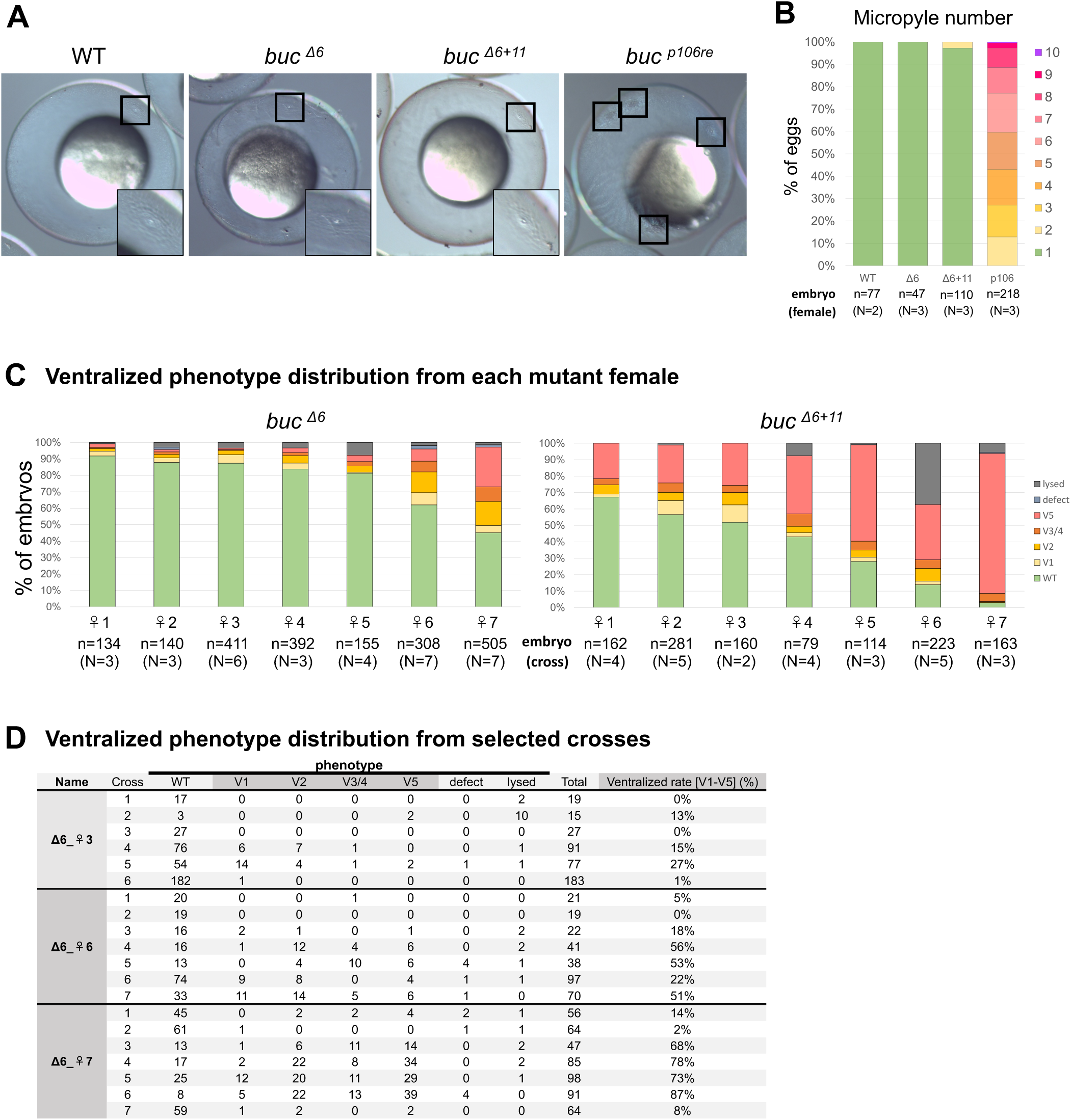
Single micropyle is formed in *buc* CRISPR mutants and ventralized phenotype is different from female to female and cross to cross. **(A)** Micropyle formation in embryos from *buc* mutant. **(B)** Micropyle number in embryos from *buc* mutant. Embryos (n) were obtained from different females (N). **(C)** Ventralized phenotype distribution from each mutant female. Embryos (n) were obtained from 7 different females from multiple crosses (N). **(D)** Ventralized phenotype distribution from selected crosses. The female’s name is consistent with that used in panel (C).

**Supplemental Figure 3, Related to figure 3 and figure 4.**
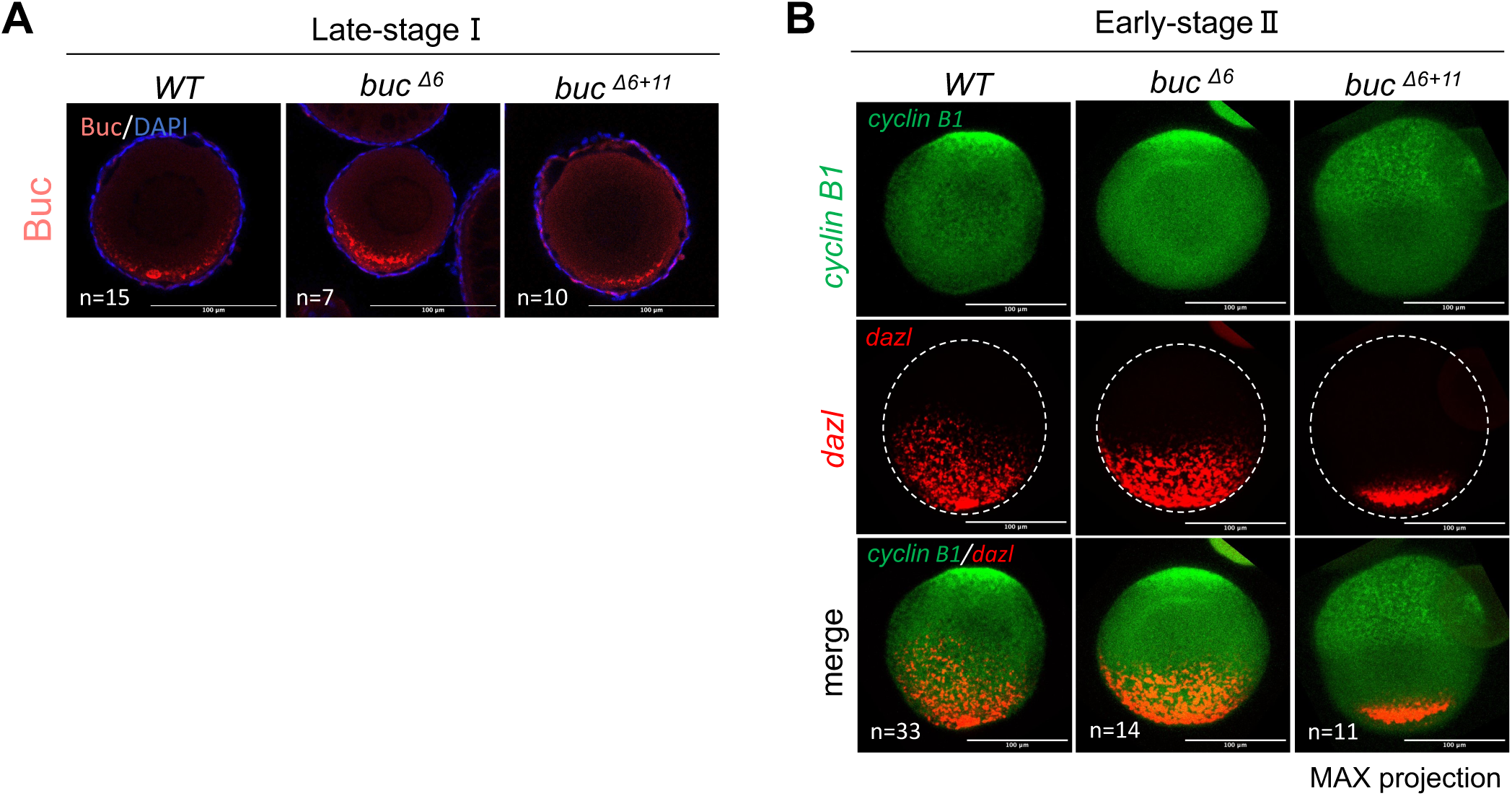
Buc protein and *dazl* mRNA are localized at the vegetal cortex in late-stage I oocyte in *buc* hypomorphic mutants. **(A)** Buc immunostaining (red) in late-stage I oocyte stained with DAPI (blue). Oocytes (n) were imaged from N=4 (WT), N=2 (*buc^Δ6^*), N=3 (*buc^Δ6+11^*) females using a confocal microscope. Scale bar = 100 μm. **(B)** Fluorescence *in situ* hybridization for *cyclin B1* (green, Animal-pole mRNA) and *dazl* mRNA (red, Vegetal-pole mRNA) in early-stage II oocyte. Z-stack images were obtained using a confocal microscope and images were generated with max intensity projection. Oocytes (n) were imaged from N=4 (WT), N=2 (*buc^Δ6^*), N=2 (*buc^Δ6+11^*) females. White dot-line outlines the oocyte. Scale bar = 100 μm.

**Supplemental Figure 4, Related to figure 6.**
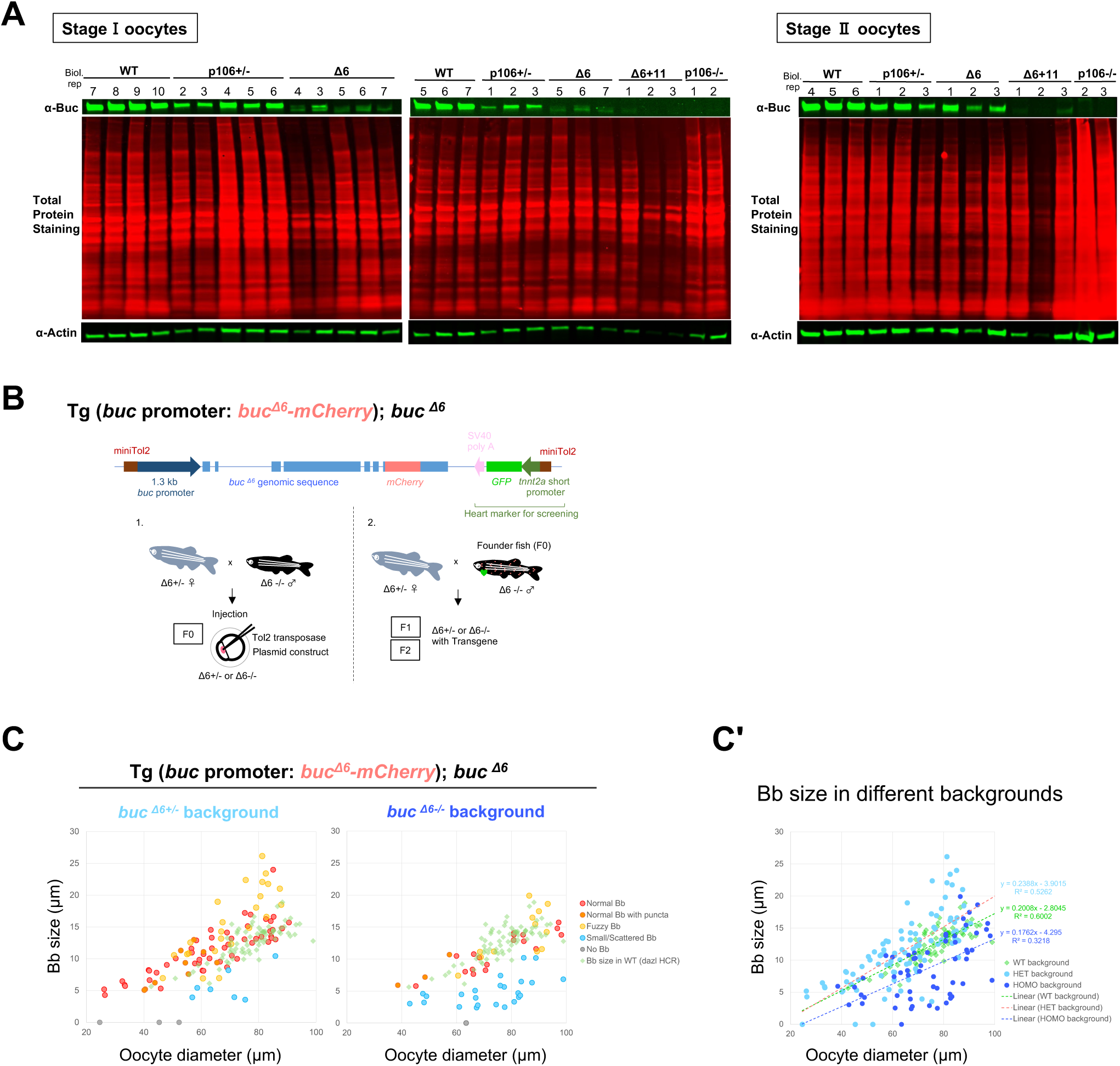
**(A)** Representative blots of total protein staining used for quantitative western blot analysis. Total protein staining was used for normalization in Figure 6C. **(B)** Establishing Tg (*buc* promoter*: buc^Δ6^-mCherry*); *buc^Δ6^* using Tol2 transposon system. **(C)** Bb size in *buc ^Δ6+/−^* or *buc ^Δ6−/−^* background Tg (*buc* promoter*: buc^Δ6^-mCherry*). Bb sizes were plotted against oocyte diameter. Bb size in WT (from *dazl* HCR, green) was plotted as a WT reference. Each dot indicates a data point from an individual oocyte, and color shows the phenotype of Bb used in Figure 6D-E. **(C’)** Bb size in different backgrounds with regression lines (green=WT, light blue=Heterozygous background, blue=Homozygous background). Analysis of covariance (ANCOVA) indicated there were significant differences in the Bb size among the backgrounds.

**Supplemental Figure 5.**
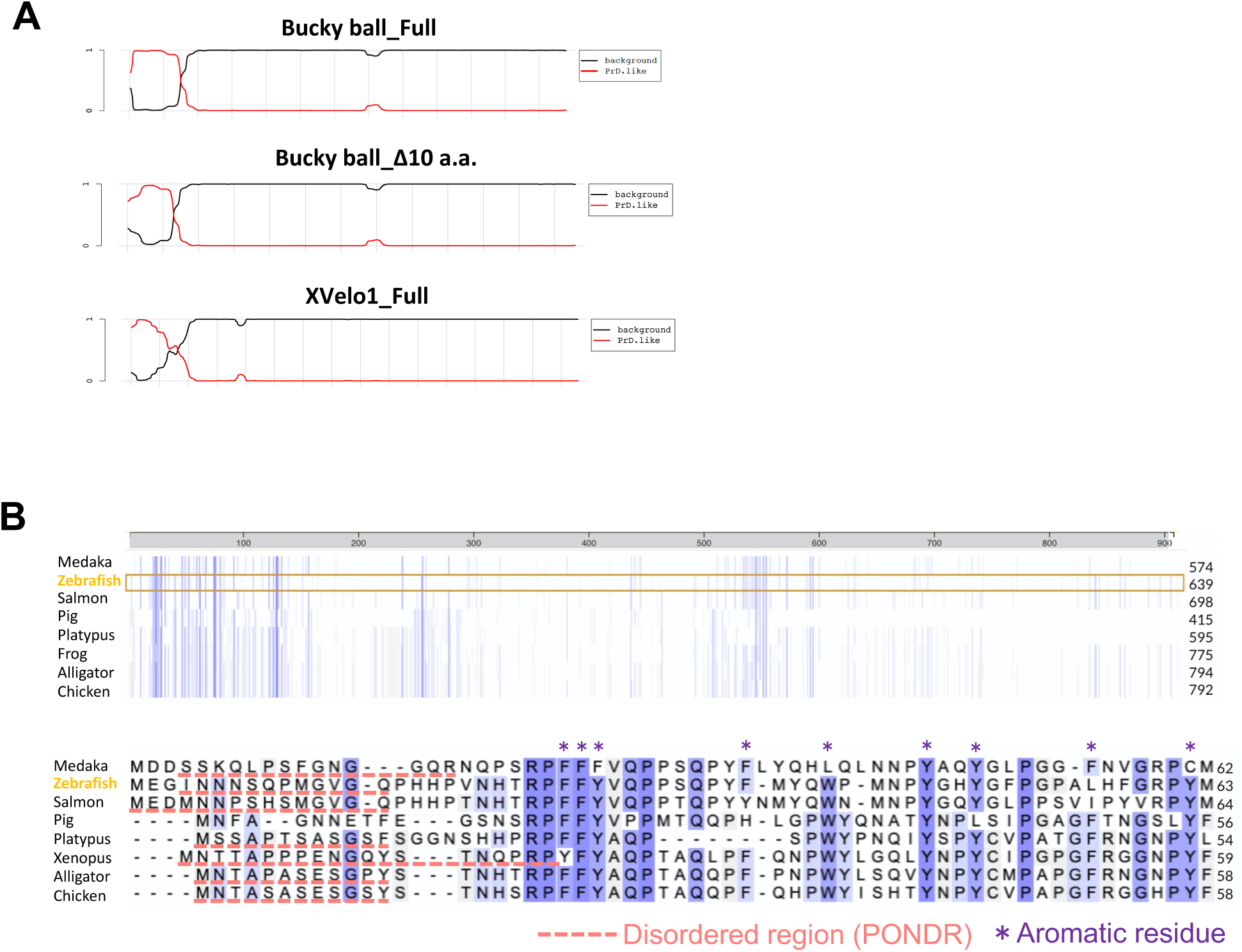
**(A)** Buc, XVelo1, and Buc with 10-amino acid deletion sequence analyzed with the prion-like domain algorithm PLAAC. Lack of the first 10 amino acids does not abolish the prion-like domain potential of the Buc protein. **(B)** Buc protein sequence comparison among species. The first 10 amino acids of Buc protein are not conserved but predicted as a disordered region by PONDR (indicated with red dot-line). Sequence alignment was performed using UniProt. An asterisk indicates the aromatic residues important for the prion-like domain.

## REFERENCES

1. Weaver, C. & Kimelman, D. Move it or lose it: Axis specification in Xenopus. Development vol. 131 3491–3499 Preprint at 10.1242/dev.01284 (2004).

2. Fuentes, R. et al. The maternal coordinate system: Molecular-genetics of embryonic axis formation and patterning in the zebrafish. in Current Topics in Developmental Biology vol. 140 341–389 (Academic Press Inc., 2020).

3. Pelliccia, J. L., Jindal, G. A. & Burdine, R. D. Gdf3 is required for robust Nodal signaling during germ layer formation and left-right patterning. Elife 6, (2017).

4. Ku, M. & Melton, D. A. Xwnt-11: a maternally expressed Xenopus wnt gene. Development 119, 1161–1173 (1993).

5. Lu, F.-I., Thisse, C. & Thisse, B. Identification and mechanism of regulation of the zebrafish dorsal determinant. Proceedings of the National Academy of Sciences 108, 15876–15880 (2011).

6. Nojima, H. et al. Syntabulin, a motor protein linker, controls dorsal determination. Development 137, 923 LP – 933 (2010).

7. Ge, X. et al. Hecate/Grip2a Acts to Reorganize the Cytoskeleton in the Symmetry-Breaking Event of Embryonic Axis Induction. PLoS Genet 10, e1004422 (2014).

8. Yan, L. et al. Maternal Huluwa dictates the embryonic body axis through b-catenin in vertebrates. Science (1979) 362, (2018).

9. Heasman, J. Maternal determinants of embryonic cell fate. Seminars in Cell and Developmental Biology vol. 17 93–98 Preprint at 10.1016/j.semcdb.2005.11.005 (2006).

10. Kloc, M., Bilinski, S. & Etkin, L. D. The Balbiani Body and Germ Cell Determinants: 150 Years Later. Curr Top Dev Biol 59, 1–36 (2004).

11. Bradley, J. T., Estridge, B. H., Kloc, M., Wolfe, K. G. & Bilinski, S. M. Balbiani bodies in cricket oocytes: Development, ultrastructure, and presence of localized RNAs. Differentiation 67, 117–127 (2001).

12. Jaglarz, M. K., Nowak, Z. & Biliński, S. M. The Balbiani body and generation of early asymmetry in the oocyte of a tiger beetle. Differentiation 71, 142–151 (2003).

13. Kosaka, K., Kawakami, K., Sakamoto, H. & Inoue, K. Spatiotemporal localization of germ plasm RNAs during zebrafish oogenesis. Mech Dev (2007) doi:10.1016/j.mod.2007.01.003.

14. Hertig, A. T. The primary human oocyte: Some observations on the fine structure of Balbiani’s vitelline body and the origin of the annulate lamellae. American Journal of Anatomy 122, 107–137 (1968).

15. Dhandapani, L. et al. Comparative analysis of vertebrates reveals that mouse primordial oocytes do not contain a Balbiani body. J Cell Sci 135, (2022).

16. Jamieson-Lucy, A. & Mullins, M. C. The vertebrate Balbiani body, germ plasm, and oocyte polarity. Curr Top Dev Biol 135, 1–34 (2019).

17. Kloc, M. & Etkin, L. D. Two distinct pathways for the localization of RNAs at the vegetal cortex in Xenopus oocytes. Development 121, 287–297 (1995).

18. Houston, D. W. Regulation of cell polarity and RNA localization in vertebrate oocytes. in International Review of Cell and Molecular Biology vol. 306 127–185 (Elsevier Inc., 2013).

19. Dosch, R. et al. Maternal control of vertebrate development before the midblastula transition: Mutants from the zebrafish I. Dev Cell 6, 771–780 (2004).

20. Marlow, F. L. & Mullins, M. C. Bucky ball functions in Balbiani body assembly and animal-vegetal polarity in the oocyte and follicle cell layer in zebrafish. Dev Biol 321, 40–50 (2008).

21. Boke, E. et al. Amyloid-like Self-Assembly of a Cellular Compartment. Cell 166, (2016).

22. Heim, A. E. et al. Oocyte polarity requires a Bucky ball-dependent feedback amplification loop. Development 141, (2014).

23. Riemer, S., Bontems, F., Krishnakumar, P., Gömann, J. & Dosch, R. A functional Bucky ball-GFP transgene visualizes germ plasm in living zebrafish. Gene Expression Patterns 18, 44–52 (2015).

24. Maegawa, S., Yasuda, K. & Inoue, K. Maternal mRNA localization of zebrafish DAZ-like gene. Mech Dev 81, 223–226 (1999).

25. Howley, C. & Ho, R. K. mRNA localization patterns in zebrafish oocytes. Mech Dev 92, 305–309 (2000).

26. Bontems, F. et al. Bucky Ball Organizes Germ Plasm Assembly in Zebrafish. Current Biology 19, 414–422 (2009).

27. Elkouby, Y. M., Jamieson-Lucy, A. & Mullins, M. C. Oocyte Polarization Is Coupled to the Chromosomal Bouquet, a Conserved Polarized Nuclear Configuration in Meiosis. PLoS Biol (2016) doi:10.1371/journal.pbio.1002335.

28. Kar, S. et al. The Balbiani body is formed by microtubule-controlled molecular condensation of Buc in early oogenesis. Current Biology 35, 315–332.e7 (2025).

29. Kozak, M. Point mutations define a sequence flanking the AUG initiator codon that modulates translation by eukaryotic ribosomes. Cell 44, 283–292 (1986).

30. Grzegorski, S. J., Chiari, E. F., Robbins, A., Kish, P. E. & Kahana, A. Natural variability of Kozak sequences correlates with function in a zebrafish model. PLoS One 9, (2014).

31. Reimão-Pinto, M. M., Castillo-Hair, S. M., Seelig, G. & Schier, A. F. The regulatory landscape of 5′ UTRs in translational control during zebrafish embryogenesis. Dev Cell 60, 1498–1515.e8 (2025).

32. Roovers, E. F. et al. Tdrd6a Regulates the Aggregation of Buc into Functional Subcellular Compartments that Drive Germ Cell Specification. Developmental Cell (2018) doi:10.1016/j.devcel.2018.07.009.

33. Yisraeli, J. K., Sokol, S. & Melton, D. A. A two-step model for the localization of maternal mRNA in Xenopus oocytes: Involvement of microtubules and microfilaments in the translocation and anchoring of Vg1 mRNA. Development 108, 289–298 (1990).

34. Zhou, Y. & King, M. Lou. RNA Transport to the Vegetal Cortex of Xenopus Oocytes. DEVELOPMENTAL BIOLOGY vol. 179 (1996).

35. Hudson, C. & Woodland, H. R. Xpat, a Gene Expressed Specifically in Germ Plasm and Primordial Germ Cells of Xenopus Laevis. (1998).

36. Allen, L., Kloc, M. & Etkin, L. D. Identification and characterization of the Xlsirt cis-acting RNA localization element. Differentiation 71, 311–321 (2003).

37. Choo, S., Heinrich, B., Betley, J. N., Chen, Z. & Deshler, J. O. Evidence for common machinery utilized by the early and late RNA localization pathways in Xenopus oocytes. Dev Biol 278, 103–117 (2005).

38. Claußen, M., Horvay, K. & Pieler, T. Evidence for overlapping, but not identical, protein machineries operating in vegetal RNA localization along early and late pathways in Xenopus oocytes. Development 131, 4263–4273 (2004).

39. Suzuki, H. et al. Vegetal localization of the maternal mRNA encoding an EDEN-BP/Bruno-like protein in zebrafish. Mech Dev 93, 205–209 (2000).

40. Lancaster, A. K., Nutter-Upham, A., Lindquist, S. & King, O. D. PLAAC: a web and command-line application to identify proteins with prion-like amino acid composition. Bioinformatics 30, 2501–2502 (2014).

41. Xue, B., Dunbrack, R. L., Williams, R. W., Dunker, A. K. & Uversky, V. N. PONDR-FIT: A meta-predictor of intrinsically disordered amino acids. Biochimica et Biophysica Acta (BBA) - Proteins and Proteomics 1804, 996–1010 (2010).

42. Putnam, A., Thomas, L. & Seydoux, G. RNA granules: functional compartments or incidental condensates? Genes & development vol. 37 354–376 Preprint at 10.1101/gad.350518.123 (2023).

43. Labun, K. et al. CHOPCHOP v3: expanding the CRISPR web toolbox beyond genome editing. Nucleic Acids Res 47, W171–W174 (2019).

44. Jing, L. Genotyping for Single Zebrafish (Fin Clip) or Zebrafish Embryo. Bio-Protocol Exchange (2012) doi:10.21769/BioProtoc.182.

45. Thisse, C. & Thisse, B. High-resolution in situ hybridization to whole-mount zebrafish embryos. Nat Protoc 3, 59–69 (2008).

46. Parichy, D. M., Elizondo, M. R., Mills, M. G., Gordon, T. N. & Engeszer, R. E. Normal table of postembryonic zebrafish development: Staging by externally visible anatomy of the living fish. Developmental Dynamics 238, 2975–3015 (2009).

47. Choi, H. M. T. et al. Third-generation in situ hybridization chain reaction: multiplexed, quantitative, sensitive, versatile, robust. Development 145, dev165753 (2018).

48. Choi, H. M. T. et al. Mapping a multiplexed zoo of mRNA expression. Development 143, 3632–3637 (2016).

49. Elkouby, Y. M. & Mullins, M. C. Methods for the analysis of early oogenesis in Zebrafish. Dev Biol 430, 310–324 (2017).

50. Zinski, J., Tuazon, F., Huang, Y., Mullins, M. & Umulis, D. Imaging and Quantification of P-Smad1/5 in Zebrafish Blastula and Gastrula Embryos. in Bone Morphogenetic Proteins: Methods and Protocols (ed. Rogers, M. B.) 135–154 (Springer New York, New York, NY, 2019). doi:10.1007/978-1-4939-8904-1_10.

51. Lee, M. T. et al. Nanog, Pou5f1 and SoxB1 activate zygotic gene expression during the maternal-to-zygotic transition. Nature 503, 360–364 (2013).

52. Jamieson-Lucy, A. & Mullins, M. C. Isolation of Zebrafish Balbiani Bodies for Proteomic Analysis. in Vertebrate Embryogenesis: Embryological, Cellular, and Genetic Methods (ed. Pelegri, F. J.) 295–302 (Springer New York, New York, NY, 2019). doi:10.1007/978-1-4939-9009-2_17.

53. Kobayashi, M., Jamieson-Lucy, A. & Mullins, M. C. Microinjection Method for Analyzing Zebrafish Early Stage Oocytes. Front Cell Dev Biol Volume 9–2021, (2021).

54. Suster, M. L., Kikuta, H., Urasaki, A., Asakawa, K. & Kawakami, K. Transgenesis in Zebrafish with the Tol2 Transposon System. in Transgenesis Techniques: Principles and Protocols (ed. Cartwright, E. J.) 41–63 (Humana Press, Totowa, NJ, 2009). doi:10.1007/978-1-60327-019-9_3.

55. Balciunas, D. et al. Harnessing a High Cargo-Capacity Transposon for Genetic Applications in Vertebrates. PLoS Genet 2, e169- (2006).

56. Kwan, K. M. et al. The Tol2kit: A multisite gateway-based construction kit for Tol2 transposon transgenesis constructs. Developmental Dynamics 236, 3088–3099 (2007).

57. Grajevskaja, V., Camerota, D., Bellipanni, G., Balciuniene, J. & Balciunas, D. Analysis of a conditional gene trap reveals that tbx5a is required for heart regeneration in zebrafish. PLoS One 13, e0197293- (2018).

58. Consortium, T. U. UniProt: the Universal Protein Knowledgebase in 2025. Nucleic Acids Res 53, D609–D617 (2025).

59. Sievers, F. et al. Fast, scalable generation of high-quality protein multiple sequence alignments using Clustal Omega. Mol Syst Biol 7, 539 (2011).

60. Schindelin, J., et al. Fiji: an open-source platform for biological-image analysis. Nat Methods 9, 676–682 (2012).

61. Gupta, T. et al. Microtubule actin crosslinking factor 1 regulates the balbiani body and animal-vegetal polarity of the zebrafish oocyte. PLoS Genet 6, 14–18 (2010).

